# Nuclear CK1δ as a Critical Determinant of PER:CRY Complex Dynamics and Circadian Period

**DOI:** 10.1101/2025.02.12.637842

**Authors:** Fidel E. Serrano, Daniela Marzoll, Bianca Ruppert, Axel C. R. Diernfellner, Michael Brunner

## Abstract

The mammalian circadian clock is governed by a feedback loop in which the transcription activator CLOCK:BMAL1 induces expression of its inhibitors, PERs and CRYs, which form a complex with CK1δ, the main circadian kinase. However, the spatiotemporal dynamics of this feedback loop and the precise role of CK1δ remain incompletely understood. Using an inducible overexpression system, we show that nuclear availability of CK1δ is limited by both rapid nuclear degradation and active export of unassembled kinase, while cytoplasmic kinase is readily available for association with PERs. We demonstrate that CK1δ-mediated phosphorylation may disrupt PER2-CRY1 interaction thereby resulting in cytoplasmic PER2 dimers containing substoichiometric amounts of CRY1. Analysis of endogenous PER2 localization in the context of an intact circadian clock reveals that PER2 accumulates in the cytoplasm late in the circadian cycle. Based on these findings, we propose that cytoplasmic accumulation of PER:CRY:CK1δ complexes contributes to the clearance of nuclear PER2, while the CK1δ-dependent release of CRY1 into the nucleus may sustain CLOCK:BMAL1 repression on DNA supporting the transition from the early to the late repressive phase.

## Introduction

Circadian clocks are molecular oscillators that regulate 24 h physiological rhythms as an adaptive response to day-night cycles^1,2^. In mammals, the core clock relies on a delayed negative transcription-translation feedback loop (TTFL), in which the transcriptional activators BMAL1 and CLOCK drive the expression of their inhibitors, PERIOD (PER) and CRYPTOCHROME (CRY) proteins^3–5^. This core loop is reinforced by a secondary feedback loop in which the nuclear receptors RORs and REV-ERBs connect the core oscillator to cellular metabolism by activating and repressing *BMAL1* transcription, respectively^6,7^. Circadian clocks are expressed in many tissues and entrained by daily recurring cues. Entraining environmental light cues are received by the suprachiasmatic nucleus (SCN), which directly and indirectly synchronizes clocks in peripheral organs via neuronal and hormonal/humoral cues. Recurring metabolic cues entrain peripheral clocks in the liver and other organs and determine their phase relative to the 24-hour light-dark cycle. The circadian system directly and indirectly controls key metabolic pathways^8–10^.

The prevailing “dual-mode of circadian repression” model offers a robust framework for understanding the circadian clock on the molecular level, distinguishing between the so-called "early" and "late" repressive phases. During the early repressive phase, a complex composed of PERs, CRYs, and casein kinase 1δ (CK1δ) assembles on CLOCK:BMAL1, leading to CLOCK phosphorylation, reducing CLOCK:BMAL1 binding to DNA, and thereby resulting in “displacement”-type repression. In the late repressive phase, phosphorylated PER proteins are degraded and the remaining CRY1 directly suppresses CLOCK:BMAL1 by sequestering the BMAL1 transactivation domain (TAD) resulting in “blocking”-type repression. Upon CLOCK dephosphorylation, the CLOCK:BMAL1 complex re-associates with DNA, establishing a poised promoter state. This configuration is characterized by transcriptional readiness, awaiting CRY1 to decrease below a threshold allowing reinitiation the circadian transcriptional cycle^11–17^. It has been shown recently that CLOCK:BMAL1 in the SCN remains partially repressed by CRY1 even at peak circadian transcription^18^, a mechanism that may ensure responsiveness of the circadian clock to synchronizing cues^18,19^.

Clock component homologs generally have redundant functions but differ in expression phases and binding affinities. PER1 and PER2 are both essential for maintaining robust circadian rhythms. PER1, expressed earlier, shortens the circadian period when PER2 is absent, whereas PER2 lengthens the period when PER1 is absent; PER3 plays a less critical role^20,21^. CRY1 and CRY2 exhibit overlapping repressive functions when complexed with PER proteins. In addition, CRY1, but not CRY2, serves as a potent repressor of CLOCK:BMAL1 after PERs are degraded^3,22,23^. As a consequence, CRY1 lengthens and CRY2 shortens circadian period^23^. The casein kinase 1 splice isoforms CK1δ1 and CK1δ2, and the similar CK1ε all support the clock, with their contributions varying potentially by tissue-specific expression^24,25^. Likewise, NPAS2 can partially substitute for CLOCK in a tissue-specific manner^26^.

PER proteins serve as limiting scaffolds for the assembly of the repressive complex, playing a key role in circadian timing. CK1δ is crucial in regulating PER stability and function by phosphorylating PER proteins at numerous sites. A central aspect of this regulation involves the phosphorylation of hPER2 at S662 (S659 in mPer2) within the FASP (Familial Advanced Sleep Phase) region by CK1δ bound to the Casein Kinase Binding Domain (CKBD)^27,28^. This priming phosphorylation triggers rapid phosphorylation of four additional equally spaced residues in the FASP region. When fully phosphorylated, the FASP region inhibits CK1δ by blocking its active site^29^. This attenuation slows the phosphorylation of hPER2 S478 (S475 in mPer2), a residue within the β-TrCP phosphodegron, thereby delaying PER2 degradation, which is thus aligned with the circadian time frame^30^. This so-called "phosphoswitch" mechanism is well-characterized, involving the phosphorylation of five residues in the FASP region and one in the β-TrCP degron of PER2^30–32^. Phosphorylation of the FASP and β-TrCP regions severely impacts circadian period length but is not essential for circadian clock function per se. PER proteins are subjected to extensive hyperphosphorylation at numerous additional sites (about 80 in PER2) distributed across the protein^33^. The kinetics of this widespread hyperphosphorylation strongly correlate with circadian timekeeping^34^, but the functional significance of these modifications remains largely unknown. While it has been shown that phosphorylation of mPER1 masks an NLS^35^, the potential impact of PER phosphorylation on subcellular localization and interaction with other proteins has not been systematically analyzed. Furthermore, CK1δ is the main circadian kinase but is implicated in other essential cellular functions including regulation of the cell cycle and Wnt signalling^36,37^. The regulation, subcellular distribution, and functional allocation of CK1δ to its various tasks have yet to be fully elucidated.

Circadian proteins are expressed at relatively low physiological levels with many studies analyzing endogenous proteins contributing to our current understanding of the clock^15,38–44^. To complement these studies and gain additional mechanistic insight, we utilized an inducible cellular overexpression approach combined with time-lapse live-cell imaging to investigate the dynamic interactions and subcellular localizations of CK1δ, PER2, and CRY1 without interference from their endogenously expressed specific binding partners. Our findings reveal that PER2 and CRY1 mutually influence their stability, as well as their nuclear-cytoplasmic shuttling dynamics and equilibrium. We demonstrate that CK1δ turnover and shuttling are modulated in a kinase activity-dependent manner such that free active nuclear CK1δ is rapidly degraded, making it a limiting factor for interactions with nuclear PER2. In contrast, CK1δ is readily available in the cytosol. Phosphorylation of PER2 by CK1δ drives, with a delay, PER2 cytosolic localization and, interestingly, disrupts PER2-CRY1 interaction. Translocation of the released CRY1 into the nucleus could then contribute to blocking-type repression of CLOCK:BMAL1.

## Results

### PER2 and CRY1 mutually affect their subcellular shuttling dynamics

To investigate the shuttling dynamics and subcellular localization of PER2, CRY1, and PER2:CRY1 complexes, we tagged PER2 at its C-terminus with mNeonGreen (PER2-mNG) and CRY1 at its N-terminus with mKate2 (mK2-CRY1) (Fig. EV1A). Proteins were expressed under the control of a doxycycline (DOX)-inducible CMV promoter in U2OStx cells harboring a TET repressor (Life Technologies, R712-07), either via transient transfection or in stable cell lines, depending on the experimental setup. This inducible overexpression strategy was deliberately chosen, as tagging and expressing the proteins at endogenous levels from their native loci would not have been suitable for addressing the questions examined in this study as specified below. PER2 contains a nuclear localization signal (NLS) and three CRM1-dependent nuclear export signals (NESs). It shuttles between the cytosol and nucleus and CRY1 facilitates its nuclear accumulation^38,40,45,46^. However, analyzing the shuttling dynamics of PER2 alone is more complex than it might initially seem, as overexpression of PER2 stabilizes and thereby supports the accumulation of endogenous CRY proteins^47,48^. Thus, analysis of PER2 localization at steady state^40^ or at late time points after transient expression^45^ likely reflect the localization of PER2:CRY complexes. We reasoned that inducing a burst of PER2 overexpression from a strong promoter would initially outcompete the cellular capacity to sufficiently accumulate endogenous CRYs. Time-resolved analysis would then allow for the characterization of PER2 shuttling dynamics before it quantitatively engages with endogenous CRYs, up to the point at which a subcellular steady state of PER2:CRY complexes is established.

In order to induce a sharp burst of PER2 overexpression, non-confluent U2OStx cells were transiently transfected with PER2-mNG. The cells were then grown for 24 h without induction in order to provide a time window for the cell cycle-dependent nuclear uptake of the transfected DNA. Subsequent DOX-induction triggered the intended sharp burst of PER2-mNG expression. The dynamics of the subcellular localization of PER2-mNG was then analyzed at 4 h intervals over a time period of 28 h by live-cell microscopy using an Incucyte (Sartorius) system. A representative time course of the subcellular shuttling dynamics of overexpressed PER2-mNG upon DOX induction is shown in Fig 1A. Four hours after DOX induction, PER2-mNG localized predominantly to the cytosol and gradually shifted to the nucleus over a period of approximately 28 hours (Fig 1A, upper panels). When CRM1-dependent nuclear export was inhibited with leptomycin B (LMB), PER2-mNG rapidly accumulated in the nucleus (Fig 1A, lower panels), indicating that PER2 alone undergoes rapid nucleocytoplasmic shuttling. Immunoblot analysis revealed that DOX-induced overexpression of either transiently transfected PER2-mNG or stably-integrated V5-PER2 led to increased levels of endogenous CRY1 (Fig 1B, C). Considering that the transfection efficiency of U2OStx cells with PER2-mNG is low (<20%), the induction of endogenous CRY1 in the transiently transfected cells was substantial. The slow nuclear accumulation of PER2-mNG correlated with expression levels of endogenous CRY1, consistent with CRY1’s role of promoting PER2 nuclear localization^46^. These data demonstrate that PER2 alone shuttles rapidly between nucleus and cytoplasm. Its predominant cytoplasmic localization prior to the accumulation of endogenous CRYs indicates that the nuclear export of PER2 is substantially faster than its nuclear import. The nuclear accumulation dynamics of PER2 correlates with the PER2-dependent accumulation of endogenous CRY1. Hence, the apparently slow export of overexpressed PER2 determined previously at steady state^40^ likely reflects the shuttling dynamics of PER2 that is in complex with endogenous CRYs.

**Fig 1.**
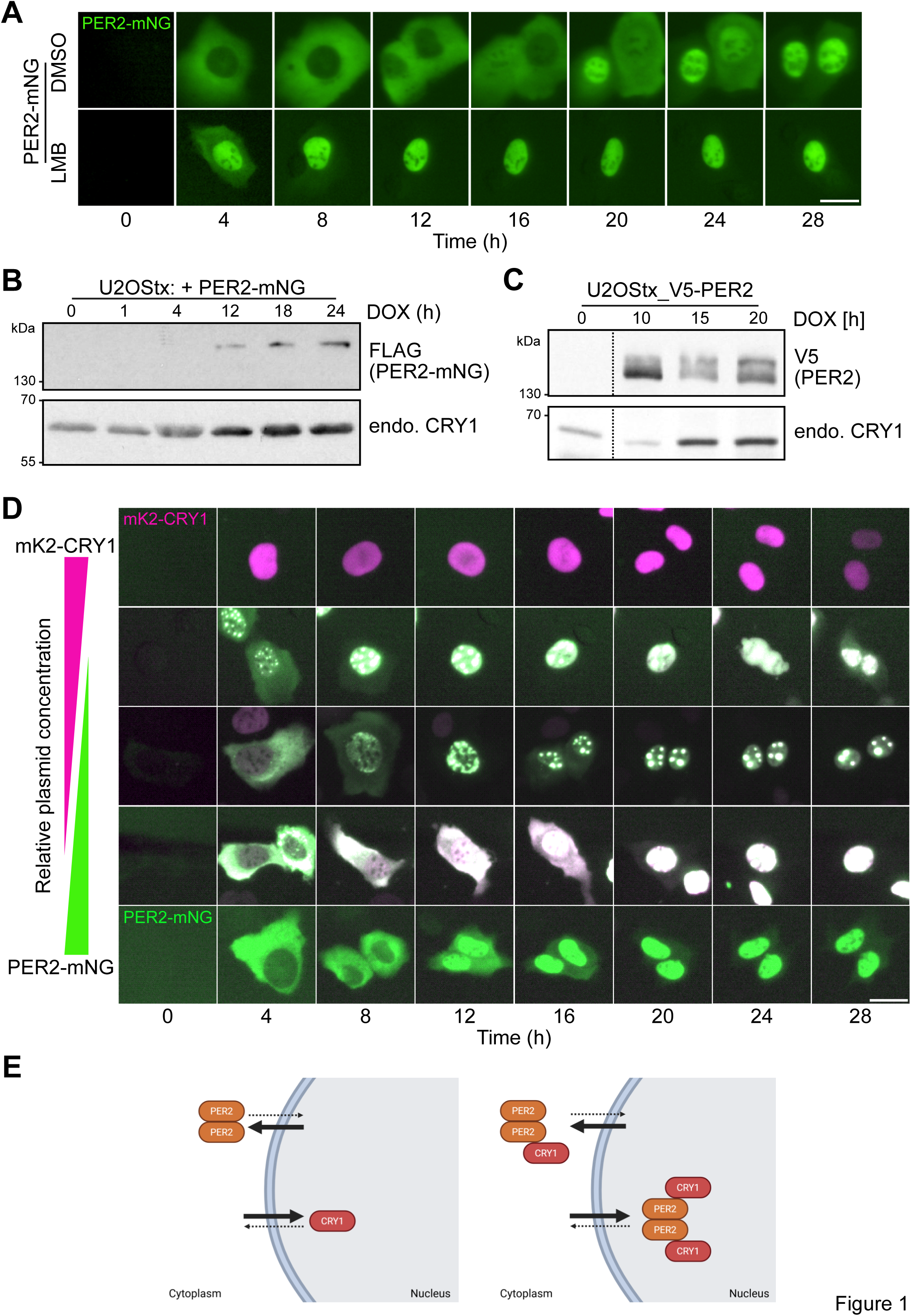
Subcellular dynamics of overexpressed PER2 and CRY1. (A) PER2-mNG exhibits rapid nucleocytoplasmic shuttling with a preference for the cytoplasm at early timepoints. DOX-induced expression of PER2-mNG in absence (DMSO) and presence of LMB. Scale bar 25 µm. n = 3. (B) Expression of PER2-mNG supports accumulation of endogenous CRY1 with kinetics correlating with nuclear accumulation of PER2-mNG. Western blot analysis of U2OStx cells transiently transfected and DOX-induced with PER2-mNG sampled at various time points post-DOX induction. (The accumulation of endogenous CRY1 in PER2-mNG-transfected cells is much higher than it appears because the transfection efficiency of U2OStx cells is <20%.) n = 3. (C) Induction of stably-integrated V5-PER2 supports accumulation of endogenous CRY1 with kinetics correlating with nuclear accumulation of PER2-mNG. Western blot analysis of V5-PER2 and CRY1 from DOX-induced U2OStx_V5-PER2 cells at the indicated time points post induction. n = 3. (D) CRY1 and PER2 mutually affect their subcellular dynamics and localization. DOX-induced transiently expressed mK2-CRY1 (top row) and PER2-mNG (bottom row) individually and in combination was analyzed over 28 h. Co-expression was assessed at mK2-CRY1 : PER2-mNG ratios of 3:1 (second row), 1:1 (third row), and 1:3 (fourth row). Scale bar 25 µm. n = 3. (E) Model of subcellular localization of PER2, CRY1, and PER2:CRY1 complexes based on their relative stoichiometries. Left: PER2 dimers, which undergo rapid nuclear–cytoplasmic shuttling, are predominantly localized in the cytoplasm, whereas CRY1 is mainly nuclear. Right: PER2 dimers bound to a single CRY1 molecule remain mainly cytoplasmic, while PER2 dimers bound to two CRY1 molecules localize to the nucleus.

To examine how PER2 and CRY1 influence each other’s subcellular dynamics, it was necessary to vary their expression levels independently while excluding contributions from endogenous PER1 and CRY2. This can only be achieved through transient transfection, as no system currently allows systematic manipulation of PER2 and CRY1 expression ratios at their endogenous loci. Although transient overexpression results in PER2 and CRY1 levels vastly exceeding those of their endogenous binding partners (i.e., PER1, CRY2, and CK1δ/ε), the overall expression levels are not high enough to saturate transient interactions with nuclear transport or degradation pathways. Therefore, U2OStx cells were transfected with PER2-mNG and mK2-CRY1 either separately or together at different ratios and, after 24 h, expression was induced with DOX. The dynamics of the subcellular localization and interaction (colocalization) of PER2-mNG and mK2-CRY1 was analyzed at 4 h intervals over a time period of 28 h (Fig 1D). Transient expression of mK2-CRY1 alone was observed after 4 h, with the protein localizing to the nucleus and remaining nuclear until it was degraded around 28 h post-DOX induction (Fig 1D, upper row). When PER2-mNG and mK2-CRY1 were coexpressed, the subcellular dynamics and expression pattern varied depending on their expression ratio (Fig 1D and EV1B). At a 1:3 transfection ratio of PER2-mNG : mK2-CRY1 (Fig 1D, 2^nd^ row), mK2-CRY1 was nuclear at all times, and nuclear accumulation of PER2-mNG was accelerated, as previously reported^46^. PER2-mNG and mK2-CRY1 accumulated in nuclear foci, which progressively decreased in number while increasing in size over time. The foci formed by the overexpressed clock proteins will be briefly characterized and discussed below. At a 1:1 transfection ratio (Fig 1D, 3^rd^ row), PER2-mNG accumulated in the nucleus more slowly, where it formed foci with mK2-CRY1. At early time points (4 h and 8 h) some mK2-CRY1 colocalized with PER2-mNG in the cytosol. At a 3:1 ratio (Fig 1D, 4^th^ row), nuclear accumulation of PER2-mNG was slow and a substantial fraction of mK2-CRY1 colocalized with PER2-mNG in the cytosol for up to 16 h. PER2-mNG by itself localized 4 h after DOX-induction predominantly in the cytosol and slowly relocalized to the nucleus over a time course of about 20 h (Fig 1D, bottom row), which correlates with the accumulation of endogenous CRYs (see Fig 1B, C). Since PER2-mNG alone accumulates in the cytosol and mK2-CRY1 alone is nuclear, the data indicate that PER2 and CRY1 mutually influence each other’s nucleocytoplasmic shuttling dynamics and subcellular distribution. PER2 forms dimers and can therefore bind either one or two CRY1 molecules. Hence, at high mK2-CRY1 levels and/or at late timepoints after DOX induction, nuclear PER2-mNG:mK2-CRY1 complexes likely represent dimers fully saturated with two mK2-CRY1 molecules. Conversely, the prolonged cytosolic presence of mK2-CRY1 at low co-transfection ratios can only be explained by partially saturated complexes, in which a PER2-mNG dimer is bound to a single mK2-CRY1. Together, these findings indicate that the relative abundance and concentration, and thus, the degree of saturation, of dimeric PER2-mNG with monomeric mK2-CRY1 governs the balance between nuclear import and export kinetics of PER2:CRY1 complexes. Given that PER2 is predominantly cytosolic and free CRY1 is primarily nuclear, and that CRY1 promotes the nuclear accumulation of PER2, our findings suggest that CRY1 can remain in the cytosol only when incorporated into asymmetric, partially saturated complexes (Fig 1E). The potential physiological relevance of such complexes will be discussed below in light of recent quantifications of cytosolic PER2 and CRY1 in SCN slices across the circadian cycle^18^.

### PERs and CRYs in nuclear foci retain their interaction specificity

The data shown above demonstrate that clock proteins have the capacity to engage in higher molecular mass assemblies. While the formation of large, thermodynamically persisting nuclear foci is clearly a consequence of overexpression, smaller assemblies may form dynamically and transiently under physiological conditions. In the context of our work, however, the foci are a convenient tool for easily analyzing the colocalization of clock proteins. To characterize the foci formed by the overexpressed clock proteins, a stable U2OStx_mK2-CRY1 line was transfected with untagged PER2 to induce formation of nuclear foci (Fig 2A). Since all cells express mK2-CRY1 and the transfection efficiency of U2OStx cells with PER2 is relatively low, the majority of cells displayed a homogenous nuclear distribution of mK2-CRY1 while fewer cells accumulated mK2-CRY1 in nuclear foci (Fig 2B and EV2A). Immunofluorescence (IF) with PER2 antibodies confirmed that overexpressed PER2 colocalized with mK2-CRY1 in nuclear foci while endogenous PER2 exhibited diffuse cellular localization in mK2-CRY1 expressing cells without foci. Further IF analysis revealed that endogenous PER1, BMAL1, and CK1δ1 colocalized in these nuclear foci (Figs 2C-E) in contrast to their dispersed subcellular distribution in cells that contained no foci (Fig EV2B-D). Notably, PER1, BMAL1 and CK1δ expression levels were elevated in cells containing nuclear PER2 and mK2-CRY1 foci compared to neighboring cells that remained untransfected with PER2-mNG (Fig 2C-E). The data suggest that overexpressed PERs and CRYs have the capacity to form nuclear foci where they maintain specific interactions with other clock proteins and support their accumulation at elevated levels. Particularly interesting was the PER2-dependent accumulation of CK1δ at elevated levels, given that little is known about how kinase expression is regulated.

**Fig 2.**
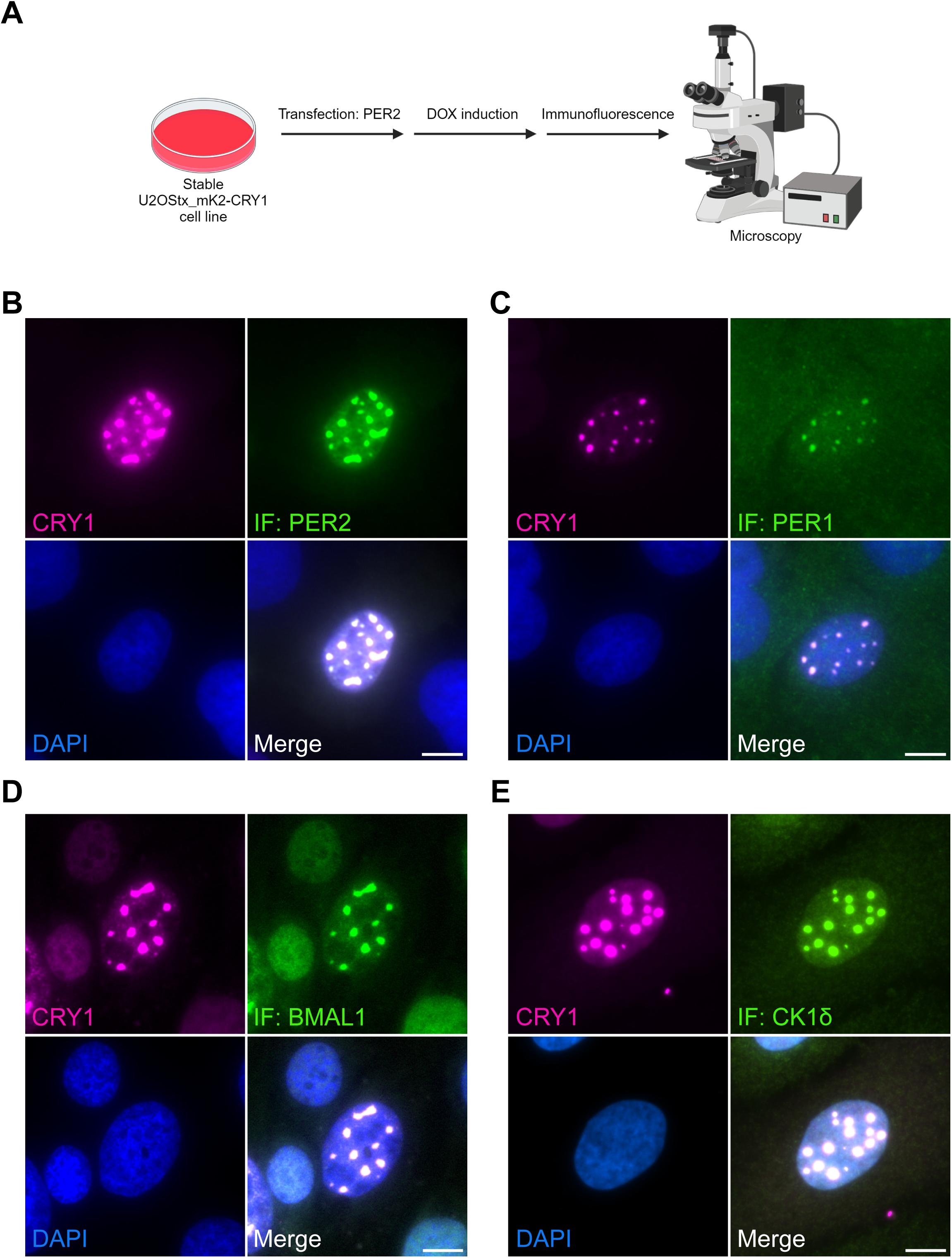
PERs and CRYs in nuclear foci retain their interaction specificity. (A) Schematic of the experimental setup: U2OStx_mK2–CRY1 cells were transfected with PER2 and induced with DOX. Cells were analyzed for mK2–CRY1 expression and by immunofluorescence (IF) using specific antibodies. (B) IF for PER2: mK2–CRY1 and PER2 co-accumulate in nuclear foci. (C) IF for PER1: nuclear foci contain endogenous PER1. (D) IF for BMAL1: nuclear foci contain endogenous BMAL1. (E) IF for CK1δ: nuclear foci contain endogenous CK1δ. Expression levels of endogenous clock proteins are elevated in transfected cells compared to surrounding untransfected cells. Scale bars: 10□µm. n = 3.

### CK1**δ** stability is regulated in an activity-dependent manner

To directly study the regulation of CK1δ and the impact of PER2 , we used previously generated stable U2OStx cell lines expressing the C-terminally FLAG-tagged wild-type CK1δ1 isoform, the tau-like mutant kinase CK1δ1-R178Q, and the kinase-dead version CK1δ1-K38R^34^. Upon DOX induction, mRNAs of the induced transgenes were expressed at similar levels, approximately 50-fold higher than the endogenous CK1δ RNA (Fig 3A). However, the proteins were expressed at vastly different levels (Fig 3B). CK1δ1 was expressed at the lowest level, CK1δ1-R178Q at an intermediate level, and CK1δ1-K38R at the highest level, suggesting an inverse correlation between resulting expression level and kinase activity or substrate specificity. Immunoblot analysis using a CK1δ-specific antibody (Fig EV3A) revealed that, in the absence of DOX, the FLAG-tagged kinases were leakily expressed at levels comparable to endogenous CK1δ (Fig 3B, bottom panel). Upon DOX induction, the FLAG-tagged kinases were overexpressed. We then tested whether the differences in resulting kinase expression were due to protein turnover. Following a cycloheximide (CHX) chase, overexpressed CK1δ1 was rapidly degraded with a half-time of less than 15 min, whereas CK1δ1-K38R and CK1δ1-R178Q remained stable (Fig 3C, E).

**Fig 3.**
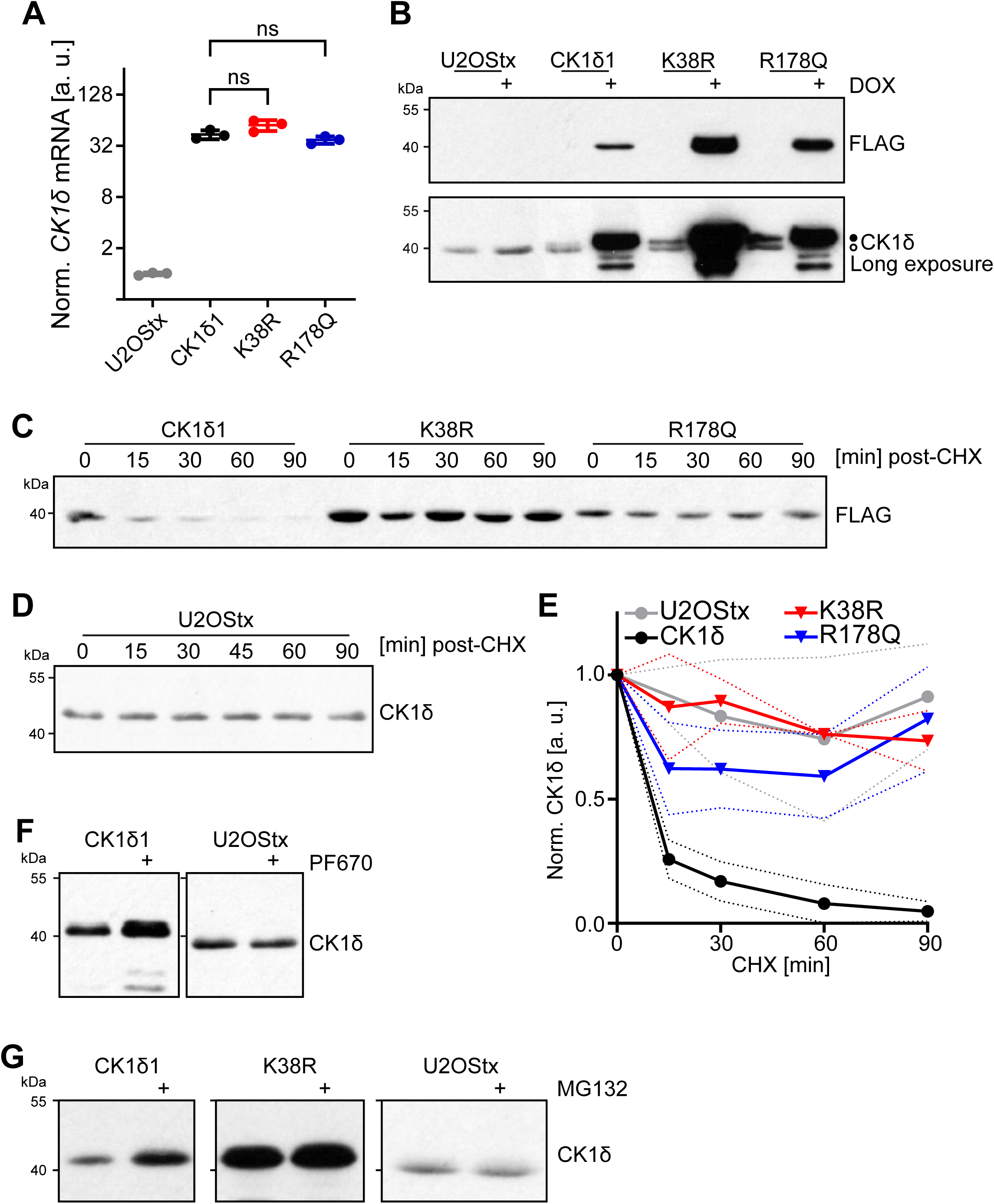
CK1δ is stability regulated in an activity-dependent manner. (A) qPCR analysis of *CK1*δ mRNA in U2OStx cells and stable U2OStx cell lines expressing DOX-inducible CK1δ1, CK1δ1-K38R, and CK1δ1-R178Q. n = 3, Students t-test, ns: not significant. n = 3. (B) Immunoblot analysis of DOX-induced CK1δ variants. Upper panel: DOX-induced expression of indicated FLAG-tagged CK1δ1 variants in stable U2OStx cell lines. Lower panel: Expression of endogenous CK1δ (open circle) and CK1δ1-FLAG variants (filled circle) analyzed with an antibody detecting CK1δ1 and CK1δ2 but not CK1ε (Figure EV2). n = 3. (C) Overexpressed CK1δ1 is unstable with a half-life of about 6 min. CK1δ1-K38R and CK1δ1-R178Q are stable over 90 min. U2OStx_CK1δ1 cells were DOX-treated for 24 h and then treated with CHX. Samples were collected after the indicated time periods (CHX chase) and analyzed by immunoblotting with FLAG antibody. (D) Endogenous CK1δ is stable over 90 min post CHX chase. (E) Quantification of (C) and (D). n = 3, solid and dashed lines: mean ± SD. (F) Left panel: Inhibition by PF670 stabilizes overexpressed FLAG-tagged CK1δ1. Right panel: 1h PF670 treatment of control cells has no effect on endogenous CK1δ levels. More protein (4x) was loaded on the gel for the analysis of endogenous CK1δ. n = 3. (G) Proteasomal inhibition by MG132 (4 h) stabilizes overexpressed (left panel) CK1δ1 and, (middle panel) to a lesser extent, the already stable CK1δ1-K38R. Right panel: MG132 has no effect on endogenous CK1δ levels in U2OStx control cells within the 4 h treatment window. n = 3.

When U2OStx control cells were treated with CHX, endogenous CK1δ remained stable over the 90 min course (Fig 3D, E), consistent with previous studies^3,11^. These data suggest that CK1δ is rapidly degraded only when overexpressed, a process facilitated by its enzymatic activity. In contrast, endogenous CK1δ, expressed at much lower physiological levels, may remain stable through interactions with various client proteins that protect it from degradation. To displace endogenous CK1δ from potential binding sites, we overexpressed CK1δ1-FLAG or mNG-CK1δ1. Overexpression of either kinase markedly reduced endogenous CK1δ levels, suggesting exchange between kinase-interacting partners and degradation of unbound kinases (Fig EV3B, C).

Since the mutant kinases CK1δ1-R178Q and CK1δ1-K38R were stable and expressed at higher levels than CK1δ1, we investigated how pharmacological inhibition with PF670462 (hereafter referred to as PF670), a well-characterized inhibitor of CK1δ^49^, affects the expression of both endogenous and overexpressed CK1δ1. Treatment with PF670 for 1 h stabilized overexpressed CK1δ1 but, consistent with previous reports^50^, did not affect the abundance of endogenous CK1δ in U2OStx control cells (Fig 3F). Our data suggest that enzymatically active, overexpressed CK1δ1 is rapidly degraded, likely because it is not assembled with client proteins. CK1δ is degraded by the proteasome via the APC/C-Cdh1 ubiquitin ligase^51^. To further characterize the degradation pathway of CK1δ, we treated cells with the proteasome inhibitor MG132 (Fig 3G). In cells overexpressing CK1δ1, a 4 h treatment with MG132 led to a significant increase in CK1δ1 abundance, indicating that, in the absence of MG132, a large fraction of CK1δ1 synthesized from overexpressed RNA was degraded. CK1δ1-K38R was also stabilized by MG132 but to a lesser extent. In contrast, MG132 treatment of U2OStx control cells did not increase endogenous CK1δ levels within the 4 h time frame.

These findings suggest that the high levels of CK1δ1 produced from abundant DOX-induced mRNA are rapidly degraded in a kinase activity-dependent manner, presumably because the kinase is not assembled with client proteins. Due to the high synthesis rate of the overexpressed CK1δ1, a 1 h inhibition of kinase activity with PF670 or a 4 h inhibition of protein turnover with MG132 was sufficient to raise CK1δ1 level to a detectable extent. In contrast, in control cells, the translation of the 50-fold less abundant endogenous CK1δ RNA (Fig 3A) was not sufficient to significantly increase kinase expression beyond the steady-state level within the inhibitor treatment time windows (Fig 3F, G). The data suggest that free CK1δ is present at low levels, potentially acting as a limiting factor for interactions with partners such as PER proteins.

### PER2 supports accumulation of endogenous CK1**δ**

To investigate whether the expression of PER2, a binding partner of CK1δ, is sufficient to elevate endogenous CK1δ levels, we generated stable U2OStx cell lines that overexpress V5-tagged PER2 in a DOX-dependent manner. Upon induction of PER2 expression, similar to endogenous CRY1 (see Fig 1B, C), endogenous CK1δ1 levels increased in parallel with PER2 requiring several hours for a detectable accumulation of the endogenous kinase (Fig 4A). The data suggest that PER2 acts as a stabilizing binding partner for newly synthesized CK1δ. A PER2-dependent increase in CK1δ1 protein levels was also observed when HEK293T cells were transfected with a constant amount of CK1δ1 plasmid and increasing amounts of PER2 (Fig 4B). In contrast, coexpression of PER2ΔCKBD, in which the CK1 binding domain (CKBD) is deleted, did not result in a substantial increase in CK1δ1 levels (Fig 4B). These findings indicate that the interaction between CK1δ1 and the CKBD of PER2 is critical for protecting the kinase from degradation. The modest stabilization of CK1δ1 observed with PER2ΔCKBD may be due to the presence of an additional, albeit weak, CK1δ1 binding site within PER2ΔCKBD^52^. Alternatively, this stabilization could occur indirectly through the PER2ΔCKBD-dependent stabilization of endogenous PERs (see Fig 1C), which could then provide a binding site for CK1δ1.

**Fig 4.**
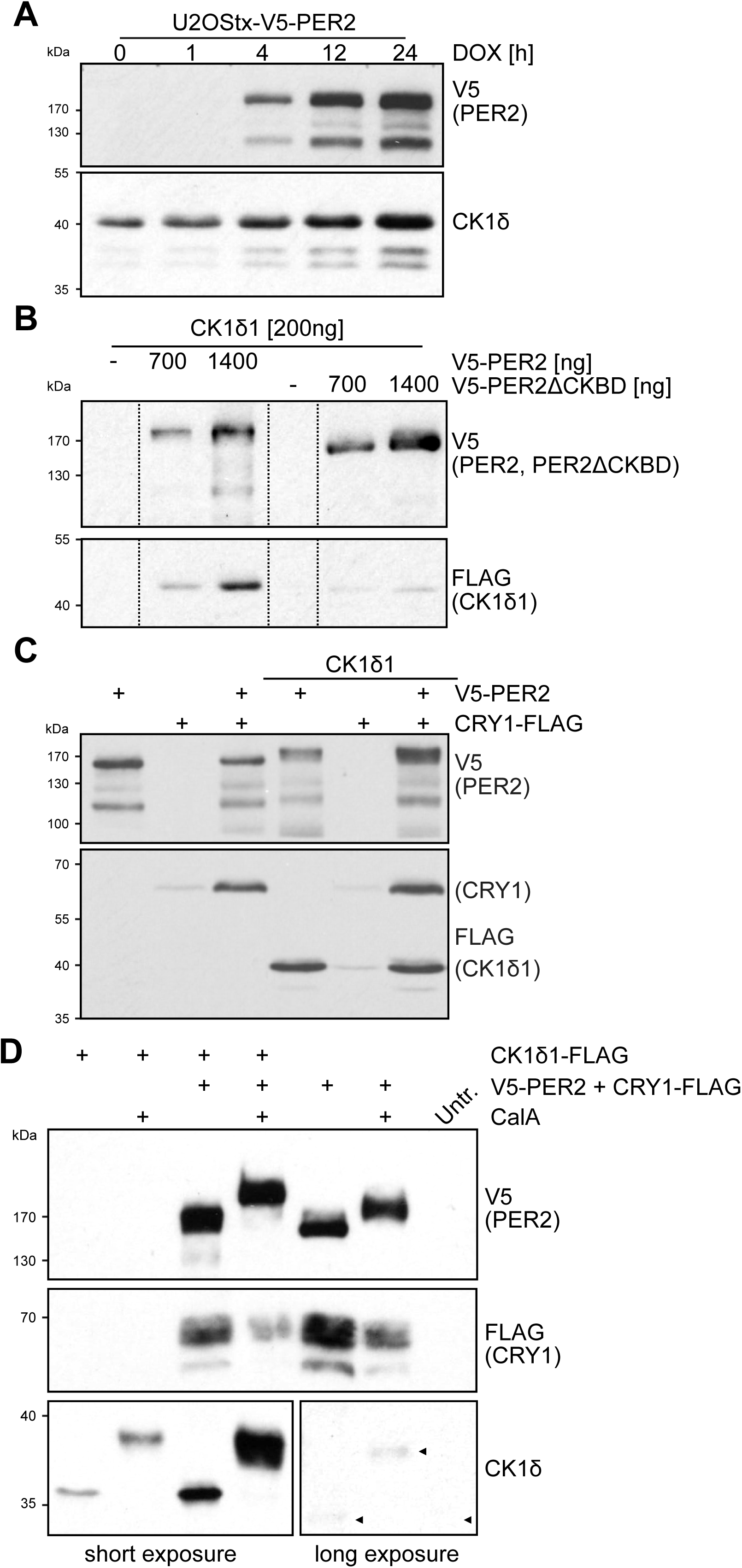
PER2 supports accumulation of endogenous CK1δ. (A) Endogenous CK1δ co-accumulates with overexpressed V5-PER2. Stable U2OStx_V5-PER2 cells were DOX-induced and analyzed after the indicated time periods for expression of (upper panel) V5-PER2 and (lower panel) endogenous CK1δ. n = 3. (B) Overexpressed CK1δ1 is stabilized dependent on PER2 dosage and PER-CK1δ interaction. HEK293T co-expression of CK1δ1-FLAG (200 ng) with increasing V5-PER2 results in V5-PER2 dose-dependent stabilization of CK1δ1-FLAG. Coexpression with V5-PER2ΔCKBD does not substantially stabilize CK1δ1-FLAG. All samples are from the same gel and Western blot with the same exposure time. Dashed vertical lines indicate that gel lanes with lower PER2 levels and size marker lanes were spliced out. n = 3. (C) PER2 stabilizes CRY1 and CK1δ1. HEK293T cell expression of indicated combinations of V5-PER2, mK2-CRY1, and CK1δ1-FLAG. n = 3. (D) Phosphatase inhibition by CalA promotes phosphorylation of PER2 as well as PER-bound CK1δ and affects levels of PER2-bound CRY1. HEK293T cell expression of indicated combinations of V5-PER2, CRY1-FLAG, and CK1δ1-FLAG with and without CalA treatment. n = 3. Upper panel: CalA promotes hyperphosphorylation of PER2. Middle panel: Levels of PER2-stabilized CRY1 correlate inversely with PER2 phosphorylation state. Lower panels: CalA promotes phosphorylation of free CK1δ1 (lanes 1 and 2) as well as of PER2-bound (stabilized) overexpressed CK1δ1 (lanes 3 and 4) and PER2-bound endogenous CK1δ (lanes 5, 6 and 7). Note, the PER2-dependent stabilization of endogenous CK1δ by comparing with untransfected HEK293T cell control.

To analyze whether CRY1 affects the stabilization of CK1δ1 by PER2, we expressed the proteins in HEK293T cells (Fig 4C). When PER2 was expressed alone, it remained hypophosphorylated. CRY1 alone was expressed at a low level. However, coexpression with PER2 led to an elevated expression of CRY1, consistent with the role of PER2 in protecting CRY1 from proteasomal degradation by occupying the FBXL3 binding site^47,48^. Coexpression of CK1δ1 and CRY1 resulted in accumulation of low levels of both proteins. In contrast, when CK1δ1 was coexpressed with PER2, whether or not CRY1 was present, CK1δ1 accumulated at a high level. Since both CRY1 and CK1δ1 were tagged with a FLAG epitope, a comparison of their expression levels indicated that PER2 stabilized both proteins to a similar extent. Given that PER2 has a dedicated binding site for CRY1 and another for CK1δ1, this is consistent with a PER2 molecule stabilizing these proteins by binding a single molecule of each. The data suggest that, under conditions of overexpression, PER2 becomes saturated with CK1δ1 and CRY1. We observed that stabilized, i.e., PER2-bound CK1δ1 was capable of hyperphosphorylating PER2 while the kinase remained hypophosphorylated itself. Upon inhibition of phosphatases with Calyculin A (CalA), both PER2 and PER2-bound CK1δ1 underwent rapid hyperphosphorylation, indicating that PER2 does not inhibit the autophosphorylation of associated CK1δ (Fig 4D). Instead, these findings suggest that cellular phosphatases act on both PER2 and PER2-bound CK1δ, with a pronounced preference for dephosphorylating the kinase. Expression of PER2 and CRY1 in HEK293T cells also led to the accumulation of endogenous CK1δ, consistent with the enrichment of CK1δ in PER2-mNG:mK2-CRY1 nuclear foci observed in U2OStx cells (see Figure 2E). However, the concentration of endogenous CK1δ remained well below the level required to saturate V5-PER2, indicating that its synthesis during the 24-hour timeframe of the experiment was insufficient to quantitatively bind all overexpressed PER2 molecules. Upon treatment with CalA, the entire population of overexpressed V5-PER2 became hyperphosphorylated, although to a lesser extent than under conditions of saturating CK1δ1. These findings suggest that PER2-anchored CK1δ, present at substoichiometric levels, phosphorylated the excess V5-PER2 in a distributive manner rather than through a fixed, obligatory 1:1 stoichiometry. Accordingly, CK1δ may dynamically equilibrate between individual PER2 molecules and/or act on multiple PER2 molecules within higher-order oligomeric assemblies. In the absence of CalA, however, the slow phosphorylation of V5-PER2 by substoichiometric levels of endogenous CK1δ was efficiently counteracted by endogenous phosphatases.

To investigate whether the stabilization of CK1δ1 by V5-PER2 was affected by kinase activity, we expressed the catalytically inactive CK1δ1-K38R and tau-like mutant kinase, CK1δ1-R178Q^34^. In the absence of V5-PER2, both mutant kinases were expressed at higher levels than CK1δ1, while CRY1 expression remained low (Fig EV4). Coexpression with PER2 further stabilized both mutant kinases, resulting in their accumulation at levels slightly higher than CK1δ1 (comp. Figs 4C and EV4 from the same blot). The data suggest that V5-PER2 stabilizes active and mutant CK1δ1 at a 1:1 ratio, while any excess kinase is degraded in a manner correlated with its kinase activity.

### Active kinase is primarily degraded in the nucleus

Active CK1δ shuttles between the cytosol and the nucleus while kinase-dead CK1δ-K38R accumulates in the nucleus^53^. To further characterize the subcellular dynamics and how it relates with kinase activity and degradation, we analyzed the localization of CK1δ, CK1δ1-R178Q and CK1δ-K38R. Overexpressed CK1δ1 was highly enriched at the centrosome and displayed some enriched accumulation in the perinuclear region, as previously described^54–56^, (Fig 5B, left panels), similar to endogenous CK1δ (Fig 5A). Kinase-dead CK1δ1-K38R accumulated in the nucleus, in agreement with a previous report^53^. It was expressed at higher levels than CK1δ1, but did not localize to the centrosome (Fig 5B, middle panels). Tau-like CK1δ1-R178Q, which is particularly compromised in its priming-dependent activity, was expressed at an intermediate level, somewhat enriched in the nucleus, and displayed centrosomal localization (Fig 5B, right panels). These results suggest that the nuclear localization of CK1δ1 and its depletion from the centrosome is inversely related to its (priming-dependent) kinase activity. We confirmed this observation by treating U2OStx_CK1δ1 cells, and U2OStx cells for control, with either MG132 or PF670. Proteasomal inhibition with MG132 for 4 h led to an increase in overexpressed CK1δ1-FLAG throughout the cell and notably also at the centrosome (Fig 5C). The data demonstrate that centrosomal binding of CK1δ was not saturated under untreated conditions, which may impact CK1δ availability for other binding partners, such as PERs. CK1δ1 inhibited for 4 h with PF670 accumulated at elevated levels (Fig 5D). As reported previously^55^ the kinase was depleted from centrosomes, and accumulated almost exclusively in the nucleus. These findings suggest that the inhibition of kinase activity affects turnover of overexpressed CK1δ1 as well as its accumulation in the nucleus and depletion from the cytosol and centrosome. Treatment of U2OStx control cells with MG132 or PF670 resulted in a small, non-significant increase in levels of endogenous CK1δ (Fig EV5A, B). These data indicate that the physiological rate of kinase synthesis in U2OStx cells is rather low and insufficient to significantly increase CK1δ expression above steady state levels within the 4 h treatment windows with MG132 and PF670.

**Fig 5.**
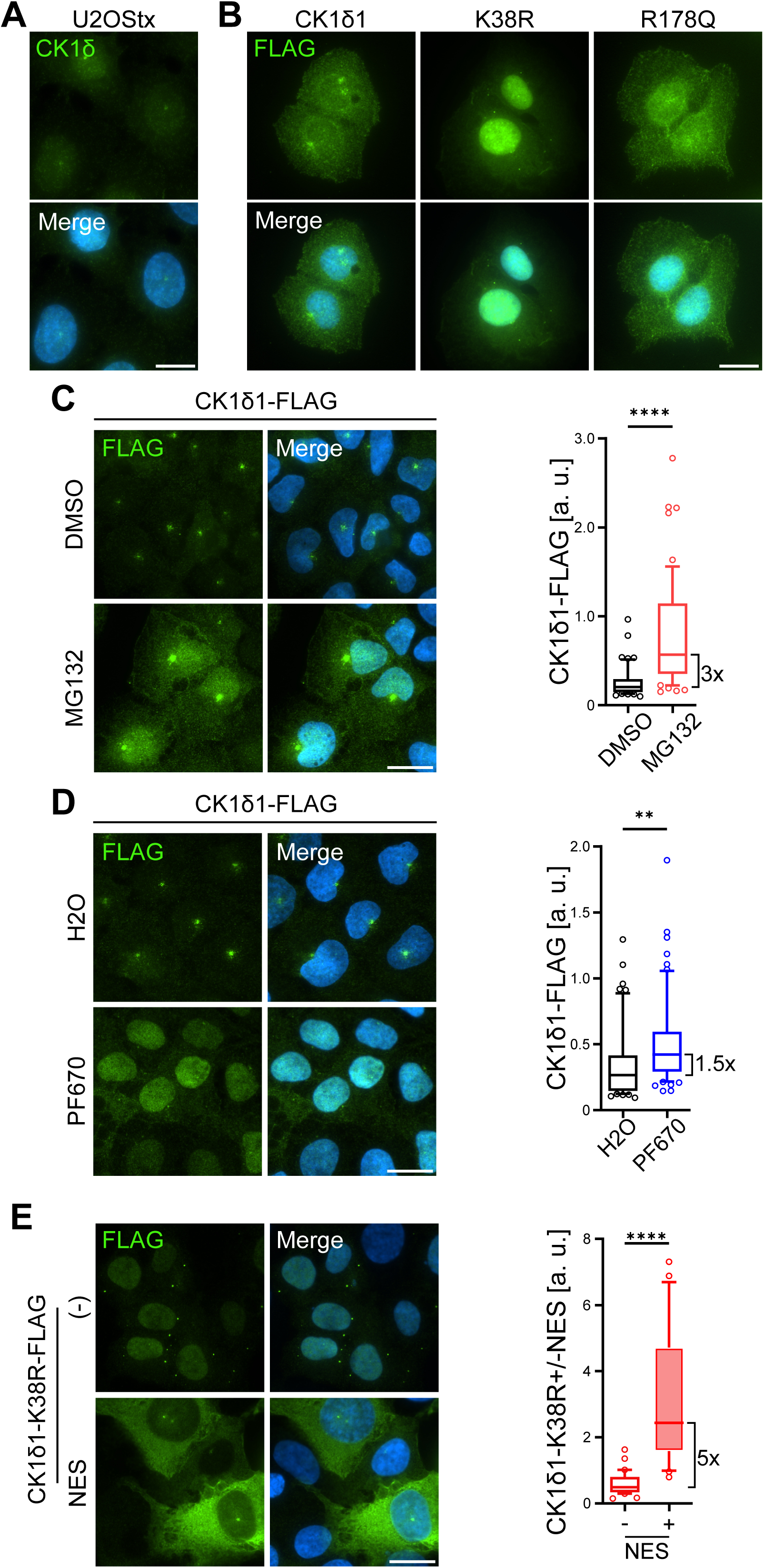
Active kinase is primarily degraded in the nucleus. (A) Endogenous CK1δ exhibits diffuse cellular localization with one distinct spot likely corresponding to the centrosome^55^. U2OStx was stained with DAPI (blue) and an antibody against CK1δ (green; AF488). Scale bar 20 µm. (B) Immunofluorescence of U2OStx cell lines overexpressing FLAG-tagged CK1δ1, CK1δ1-K38R and CK1δ1-R178Q. Cells were DOX-treated for 24 h and stained via FLAG antibody (green; AF488) and with DAPI (blue). Scale bar 20 µm. n = 3. (C) Proteasomal inhibition by MG132 results in accumulation of overexpressed CK1δ1 primarily in nucleus. Stable U2OStx_CK1δ1 cells were DOX-induced for 24 h and treated with 5 µM MG132 for 4 h. Left: IF. Right: Quantification of CK1δ1 levels in MG132-treated cells versus vehicle (DMSO) control. n ≥ 50 cells; mean ± SD; ****: p < 0.0001. Scale bar 20 µm. (D) Kinase inhibition results in accumulation of overexpressed FLAG-tagged CK1δ specifically in the nucleus. Stable U2OStx_CK1δ cells were DOX induced for 24 h and treated with 1 µM PF670 for 1 h. Left: IF. Right: Quantification of CK1δ1 levels in PF670-treated cells versus vehicle (H_2_O) control. n ≥ 50 cells; mean ± SD; **: p < 0.01. Scale bar 20 µm. (E) NES-tagged FLAG-CK1δ1-K38R accumulates at elevated levels in the cytoplasm and localizes to the centrosome. U2OStx cells were transiently transfected with either FLAG-CK1δ-K38R or NES-FLAG-CK1δ1-K38R. Left: FLAG-IF. Right: Quantification of kinase levels. n ≥ 20 cells; mean ± SD; ****: p < 0.0001. Scale bar 20 µm.

To characterize CK1δ1 degradation independently of kinase activity we transiently transfected U2OStx cells with either CK1δ1-K38R-FLAG or CK1δ1-K38R-FLAG containing an N-terminal HIV-1 nuclear export signal (NES)^57^. While CK1δ1-K38R localized to the nucleus and was depleted from the centrosome, NES-CK1δ1-K38R was found in the cytosol and the centrosome (Fig 5E). These results demonstrate that CK1δ1 facilitates nuclear export but not its localization to the centrosome per se. NES-CK1δ1-K38R was expressed at an ∼5-fold higher level than CK1δ1-K38R (Fig 5E), suggesting that inactive kinase, despite its increased stability compared to CK1δ1, is preferentially degraded in the nucleus. Overall, the data indicate that unassembled active CK1δ1 is highly unstable and preferentially degraded in the nucleus (Fig EV5C). The enzymatic activity of CK1δ1 promotes both its nuclear export and the accelerated degradation of unassembled CK1δ1 in the nucleus, potentially through a shared mechanism such as activity-dependent release from nuclear binding sites.

### CK1**δ** facilitates cytoplasmic accumulation of PER2 and release of CRY1

To analyze whether CK1δ activity affects the localization and interaction of PER2 and CRY1, we transfected U2OStx_CK1δ1 and U2OStx_CK1δ1-K38R stable cells with PER2-mNG and mK2-CRY1. Expression and localization of the DOX-induced proteins was analyzed by live cell microscopy over a time course of 28 h. In U2OStx_CK1δ1-K38R, PER2-mNG alone accumulated initially in the cytosol and relocated slowly to the nucleus (Fig EV6A, upper panels), similar to what was observed in absence of kinase (see Fig 1D). However, in U2OStx_CK1δ1 cells, PER2-mNG remained cytosolic (Fig EV6A, lower panels). Considering that PER2-mNG shuttles rapidly between the nucleus and cytosol (see Fig 1A), the data suggest that co-expression of active CK1δ1 promotes cytoplasmic accumulation of PER2, potentially through inactivation of the NLS as shown for PER1^35^. Upon expression of PER2 alone or coexpression of PER2-mNG and mK2-CRY1 in U2OStx_CK1δ1-K38R cells, PER2:CRY1 complexes accumulated in the nucleus and formed foci, which persisted for 28 h and longer (Fig EV6A and Fig 6A, upper panels), similar to what was observed in absence of overexpressed kinase (see Fig 1D, middle panels). The data indicate that the catalytically inactive CK1δ1-K38R did not affect the expression and localization of PER2-mNG and mK2-CRY1. Notably, PER2-mNG and mK2-CRY1 levels decreased in parallel over the 28-hour course of the experiment, with no free mK2-CRY1 remaining in the nucleus. In contrast, in U2OStx_CK1δ1 cells, PER2-mNG alone remained dispersed in the cytosol (Fig EV6A, lower panels), while PER2-mNG and mK2-CRY1 accumulated in cytosolic foci with a substantial fraction of mK2-CRY1 accumulating in the nucleus independently of PER2-mNG (Fig 6A, lower panels and EV6B). The foci grew in size up to about 12-16 h post DOX-induction. After approximately 20 hours, PER2-mNG was degraded, while mK2-CRY1 remained expressed and was exclusively detected in the nucleus. Nuclear mK2-CRY1 levels declined over time, but with substantially slower kinetics than those observed for cytoplasmic foci containing mK2-CRY1 and PER2-mNG. This suggests that free nuclear mK2-CRY1 represents a pool that either migrates to the nucleus upon phosphorylation-dependent PER2-mNG degradation and/or is released from PER2-mNG, potentially regulated by phosphorylation.

**Fig 6.**
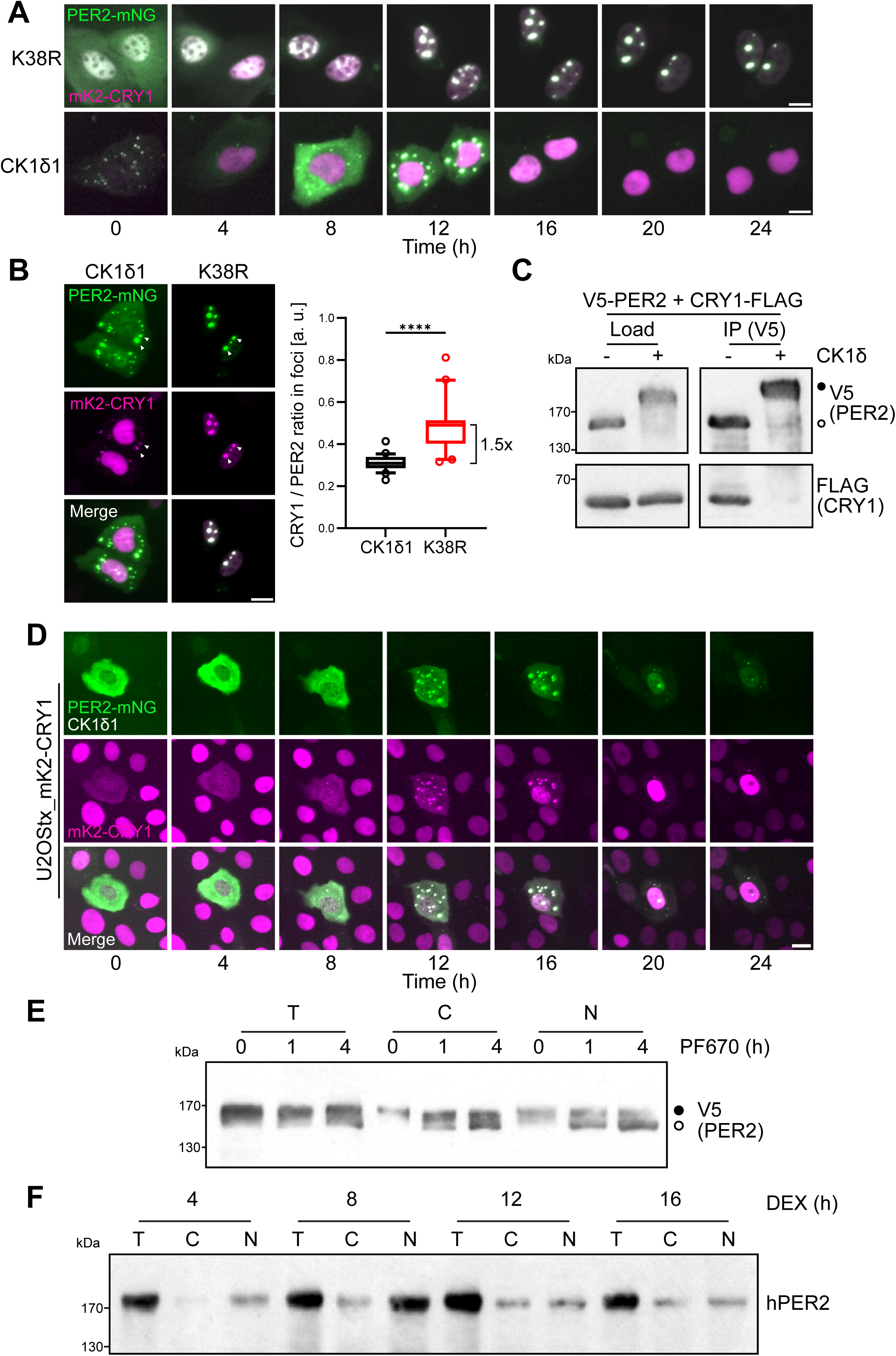
CK1δ facilitates cytoplasmic accumulation of PER2 and release of CRY1. (A) Transient transfection and coexpression of PER2-mNG and mK2-CRY1 in U2OStx_CK1δ1 and U2OStx_CK1δ1-K38R stable cell lines. Expression was DOX-induced 24 h post transfection and cells were then analyzed for 24 h with an Incucyte system. PER2-mNG and mK2-CRY1 accumulate in nuclear foci when coexpressed with inactive CK1δ1-K38R (upper row) and in cytoplasmic foci when coexpressed with active CK1δ1 (lower row). Scale bar 20 µm. (B) Cytoplasmic PER2-mNG foci (CK1δ1 cells) contain less mK2-CRY1 than nuclear foci (CK1δ1-K38R cells). Scale bar 20 µm. Right: Quantification of CRY1 : PER2 ratio in cytoplasmic and nuclear foci. n > 30 foci, mean ± SD; ****: p < 0.0001. (C) PER2 phosphorylation attenuates interaction with CRY1. V5-PER2 and CRY1-FLAG were transiently expressed in HEK293T. Cell lysate was prepared and incubated with and without recombinant CK1δΔtail for 4 h. V5-PER was then immunoprecipitated with V5 beads. Lysate (Input) and an 8x equivalent of the IP were analyzed by Western-blot. FLAG-CRY1 co-immunoprecipitated with hypophosphorylated but not with hyperphosphorylated V5-PER2. n = 3. (D) Cytoplasmic PER:CRY foci persistently release CRY1 into the nucleus in a CK1δ1-dependent fashion. Scale bar 20 µm. Central cell: Representative U2OStx_mK2-CRY1 cell co-transfected with PER2-mNG and CK1δ1, resulting in the formation of cytoplasmic PER:CRY foci. Over time, PER2-mNG levels decrease, while mK2-CRY1 accumulates in the nucleus and remains at persistently high levels, even 36 hours post DOX induction (see Fig EV6 for more examples). Surrounding cells: U2OStx_mK2-CRY1 cells not transfected with PER2-mNG. In these cells, mK2-CRY1 accumulates in the nucleus initially, but its levels decrease over time, likely due to DOX metabolism, with low levels observed 36 hours post DOX induction. (E) CK1δ inhibition by PF670 facilitated the nuclear accumulation of hypophosphorylated PER2. V5-tagged PER2 was transiently expressed in HEK293T cells together with FLAG-tagged CRY1 and CK1δ1. PF670 was added, and the subcellular localization of PER2 was analyzed at the indicated time points following PF670 treatment. (F) Endogenous PER2 subcellular redistribution from the nucleus to the cytosol in U2OStx cells was analyzed over a circadian cycle. U2OStx cells were synchronized with DEX at t = 0 h. After 4 and 8 h, endogenous PER2 was enriched in the nucleus. At later timepoints, a fraction of PER2 was localized in the cytosol.

We then quantified the relative ratio of mK2-CRY1 to PER2-mNG in cytosolic versus nuclear foci. This analysis revealed that the cytoplasmic PER:CRY foci in cells expressing active CK1δ1 contained less mK2-CRY1 compared to the nuclear PER:CRY foci in cells expressing the inactive CK1δ1-K38R (Fig 6B), suggesting that a fraction of mK2-CRY1 is released from cytoplasmic foci prior to degradation of PER2-mNG. To investigate whether CK1δ1-mediated phosphorylation triggers the release of CRY1 from PER2 independently of phosphorylation-dependent PER2 degradation, we prepared a native cell extract from HEK293T cells overexpressing V5-PER2 and CRY1-FLAG which was then incubated with purified CK1δΔtail in an *in vitro* phosphorylation reaction^34^. In the presence of kinase, PER2 was hyperphosphorylated while the electrophoretic mobility of CRY1 did not shift (Fig 6C). Subsequent immunoprecipitation revealed that CRY1 co-immunoprecipitated with unphosphorylated but not with hyperphosphorylated V5-PER2 (Fig 6C). The data suggest that CK1δ1-mediated phosphorylation of PER2 promotes the release of bound CRY1. Since free CRY is less stable, this result is consistent with the data shown above (Figure□4D, middle panel), where CalA-induced hyperphosphorylation of PER2 resulted in reduced CRY1 levels. To explore this further, we transfected stable U2OStx_mK2-CRY1 cells with PER2-mNG and untagged CK1δ1, and induced expression with DOX (Fig 6D and EV6C). The majority of cells expressed only mK2-CRY1, indicating that they remained untransfected. In contrast, cells exhibiting cytoplasmic foci of PER2-mNG were identified as cotransfected with untagged CK1δ1, as the foci become cytoplasmic only under these conditions. Tracking the DOX-induced protein dynamics over a 28-hour time course, we observed that mK2-CRY1 was depleted in untransfected cells, likely due to metabolism of DOX^58^ and degradation of previously synthesized mK2-CRY1. Interestingly, in transfected cells, mK2-CRY1 initially colocalized with PER2-mNG in cytoplasmic foci. Upon PER2-mNG degradation, liberated mK2-CRY1 translocated and accumulated at elevated levels in the nucleus. This led to sustained nuclear levels of mK2-CRY1 in cells initially expressing PER2-mNG and CK1δ1, even at later time points when mK2-CRY1 had already been depleted in neighbouring untransfected U2OStx_mK2-CRY1 cells.

To investigate the impact of phosphorylation on PER2 subcellular localization, we co-expressed V5-PER2, FLAG-CRY1, and FLAG-CK1δ in HEK293T cells. The cells were then treated with the CK1 inhibitor PF670, and the subcellular localization of PER2 was analyzed. Before PF670 treatment, PER2 was hyperphosphorylated and predominantly localized in the cytosol. Treatment with PF670 for 1 and 4 hours led to dephosphorylation indicating that PER2 phosphorylation was antagonized by phosphatases. This dephosphorylation facilitated nuclear accumulation of a substantial fraction of PER2 (Fig 6E). Finally, to assess the impact of PER2 phosphorylation on its subcellular localization within the context of a functional circadian clock, PER2 localization was monitored over a circadian cycle in dexamethasone (DEX)-synchronized U2OS cells (Fig 6F and EV6D). At early time points after synchronization, PER2 was predominantly localized in the nucleus but partially redistributed to the cytosol at later time points, consistent with previous microscopy and immunoblot analyses of the subcellular distribution of endogenously tagged PER2^15,43,59^.

Cytoplasmic localization of PER2 and CRY1 has also been reported under physiological conditions in the SCN^18^. Consistent with our observations, the CRY1-to-PER2 ratio in SCN neurons was significantly lower in the cytosol than in the nucleus. It has previously been shown that nuclear CRY1 forms a repressive complex with CLOCK:BMAL1 late in the circadian cycle^11,15^. Our data suggest that cytoplasmic PER2, and the resulting cytoplasmic PER2:CRY1 complexes, may function as a reservoir that gradually releases CRY1 into the nucleus in a CK1δ1-dependent manner, a process formally analogous to the de novo synthesis of CRY1 from its mRNA. In this model, phosphorylation-dependent cytoplasmic retention of PER2 and the slow release of CRY1 would serve to decouple CRY1’s subcellular localization, repressive activity, and degradation from those of PER2.

### Nuclear CK1**δ**1 affects circadian period length

Our data show that overexpressed and unassembled CK1δ1 in the nucleus is rapidly degraded or exported. To analyze how nuclear and cytosolic CK1δ1 affect the circadian clock, we constructed V5-CK1δ1 fusion proteins with an N-terminal mNeonGreen tag followed by either an SV40 nuclear localization signal (NLS)^60^ or an HIV-1 nuclear export signal (NES)^57^ (Fig 7A). To characterize the subcellular localization and stability of the kinases, U2OStx cells were transiently transfected with these mNG-CK1δ1 constructs. Next day, the expression was induced with DOX for 24 h, and cells were then treated with or without MG132 for 4 h. mNG-NLS-CK1δ1 was expressed at a low level and localized to the nucleus (Fig 7b, left panel), while mNG-NES-CK1δ1 accumulated in the cytosol (Fig 7b, right panel), demonstrating that the added localization signals determined the subcellular distribution of the shuttling kinases. MG132 treatment stabilized mNG-NLS-CK1δ1 about 8-fold (Fig 7B), while mNG-NES-CK1δ1 was stabilized only about 1.8-fold (Fig 7C). The data indicate that, in the absence of MG132, overexpressed mNG-NLS-CK1δ1 is rapidly degraded by the nuclear proteasome, while overexpressed mNG-NES-CK1δ1 is turned over at a lower rate because it escapes efficient nuclear degradation by enhanced export into the cytosol. We then generated and selected stable U2OStx cell lines expressing in DOX-inducible manner similar levels of mNG-NLS-CK1δ1 and mNG-NES-CK1δ1, respectively (Fig 7D). To monitor the circadian rhythm, cells were transfected with a *Bmal1* promoter (*Bmal1p*)-*luc* reporter. Cells were then incubated with or without DOX and *Bmal1p-luc* expression was recorded for 96 h. Expression of mNG-NLS-CK1δ1 shortened the circadian period by 4 h (Fig 7E, upper panel and F), while mNG-NES-CK1δ1 had no impact on period length (Fig 7E, lower panel and F). The data demonstrate that cytoplasmic mNG-NES-CK1δ1 does not interfere with the function of endogenous PER proteins, indicating that CK1δ is not functionally limiting in the cytosol. Because period length is determined by nuclear CK1δ activity, these findings further imply that the nuclear CK1δ pool is insufficient to fully saturate PER proteins, regardless of where newly synthesized PERs initially acquire the kinase (Fig. EV7).

**Fig 7.**
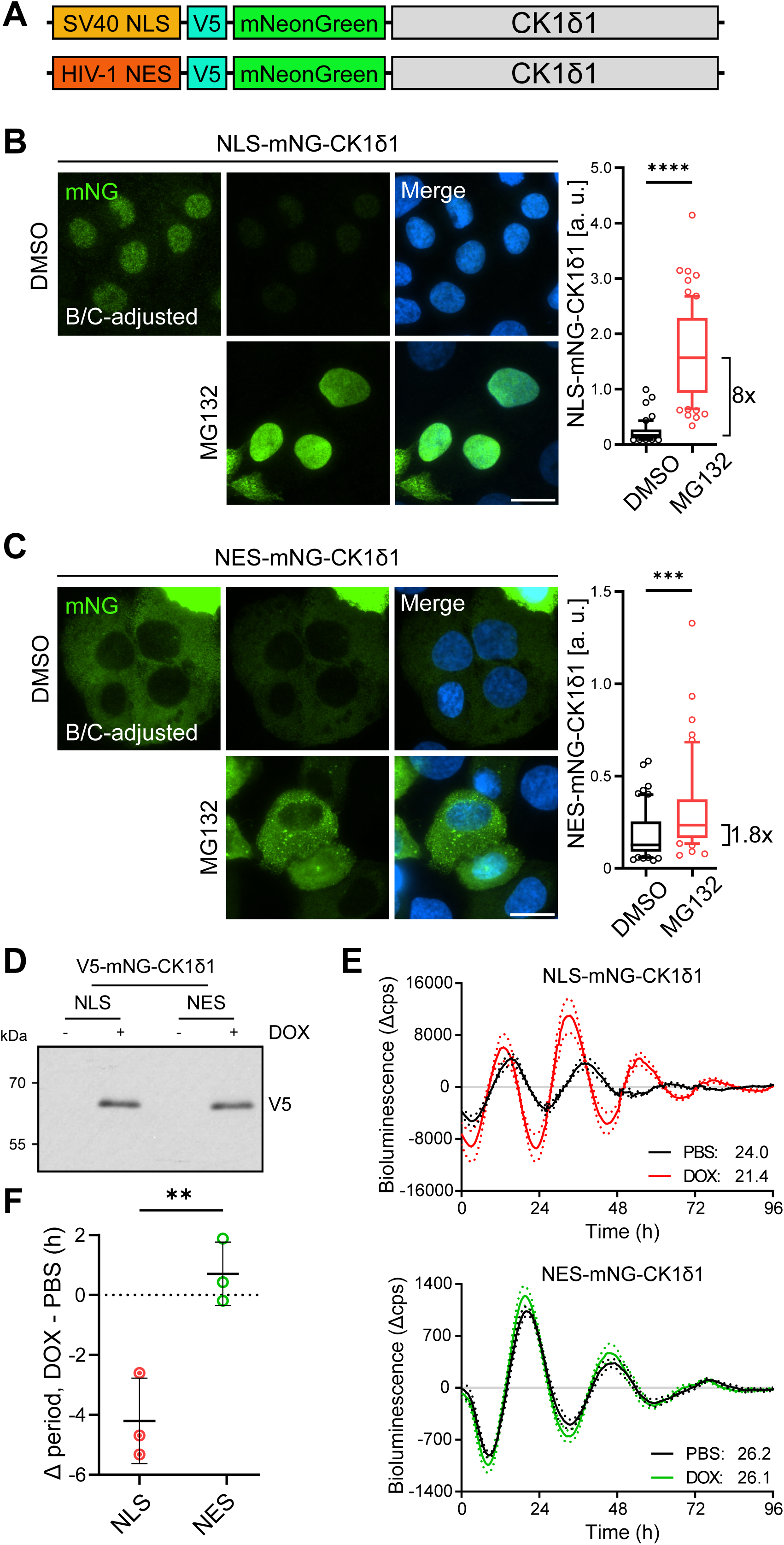
Nuclear CK1δ affects circadian period length. (A) Schematic of mNG-V5-NLS-CK1δ1 and mNG-V5-NES-CK1δ1 constructs NLS: SV40^57^; NES: HIV-1^60^. (B) Left: mNG-NLS-CK1δ1 is expressed at a very low level in the nucleus and is stabilized by MG132. B/C-adjusted panel: brightness and contrast adjusted to better visualize the kinase. Right: quantification. n > 30 cells, mean ± SD; ****: p < 0.0001. Scale bar 20 µm. (C) Left: mNG-NES-CK1δ localizes to the cytosol and is stabilized by MG132. Right: quantification. n > 30 cells, mean ± SD; ***: p < 0.001. Scale bar 20 µm. (D) Stable U2OStx lines expressing comparable levels of DOX-induced V5-tagged mNG-NLS-CK1δ1 or mNG-NES-CK1δ1. n = 3. (E) DOX-induced expression of NLS-CK1δ1 results in significant period shortening whereas NES-CK1δ1 has no effect on period. *Bmal1* promoter-luciferase assay. Representative bioluminescence traces are shown as a detrended average of 4 technical replicates of PBS- and DOX-treated cells. Resulting period was calculated using cosinor analysis built-in GraphPad Prism 10. n = 3. (F) Circadian period change comparing DOX- vs. PBS-treated mNG-NLS-CK1δ1 or mNG-NES-CK1δ1 cell lines. n = 3, mean ± SD; **: p < 0.01.

## Discussion

The current dual-mode circadian repression model describes three distinct phases of CLOCK:BMAL1 activity, regulated in temporal succession by CRY1/2, PER1/2, and CK1δ^13–17,59^. Initially, CLOCK:BMAL1 is predominantly repressed via PER:CRY:CK1δ-mediated displacement from its target genes, followed later by CRY1-dependent blocking of DNA-bound CLOCK:BMAL1, and finally derepression of CLOCK:BMAL1 when PER proteins are degraded and CRY1 levels decline below a critical threshold. Within the framework of current circadian clock models, CK1δ is assumed to be non-limiting and readily available to PER proteins as they are synthesized in the cytosol, thereby ensuring robust and precise timekeeping.

The data presented here suggest that freely available CK1δ is limited within the nucleus. Previous work has shown that nuclear export of CK1δ depends on its kinase activity^53^. We confirm these findings and extend them by demonstrating that unassembled nuclear CK1δ is subject to rapid degradation, with turnover further accelerated by its catalytic activity. Mechanistically, the activity dependence of both processes may reflect a shared mechanism, such as activity-dependent release from nuclear binding sites. Moreover, we show that CK1δ is stabilized through association with PER2. The data align with earlier findings and provide a mechanistic explanation for why nuclear CK1δ levels in mouse liver remain very low in the absence of PER proteins, but increase in parallel with the circadian accumulation of nuclear PER, while cytoplasmic CK1δ levels remain constant and higher than those in the nucleus^59^. Complex formation between CK1δ and PER proteins is well established as essential for circadian timekeeping^11^, with CK1δ1 overexpression known to shorten circadian period length^34^. Our results suggest that the subcellular distribution of CK1δ is finely tuned and suggest that nuclear concentrations remain below the threshold required for PER saturation. Specifically, overexpression of NLS-tagged CK1δ shortens period length, whereas NES-tagged CK1δ has no detectable effect. This is consistent with previous studies showing that PER associates with CK1δ in the cytosol and enters the nucleus facilitated by CRY^3,15,59^. Because each PER molecule can import only one kinase, a process that likely occurs at wild-type levels of cytosolic CK1δ, overexpression of NES-tagged CK1δ does not increase the nuclear CK1δ pool. When the few CK1δ molecules co-imported with PER:CRY complexes equilibrate with the unbound nuclear pool, rebinding to PER is constrained by rapid degradation of nuclear CK1δ or its export to the cytosol. In contrast, NLS-tagged CK1δ overexpression directly increases nuclear kinase abundance by antagonizing nuclear degradation as well as export.

Thus, when evaluating the impact of CK1δ/ε mutants on the circadian clock, not only kinase activity but also protein stability and subcellular distribution must be taken into account. Indeed, we observed that the altered activity of the tau-like CK1δ1-R178Q mutant affects its subcellular localization and turnover, leading to its accumulation at elevated levels in the nucleus. Analysis of the PER2 phosphoswitch provides strong evidence that the tau mutation (e.g., R178C in CK1δ) shortens the circadian period by attenuating phosphorylation of the FASP region in PER2 while accelerating phosphorylation of the β-TrCP degron site (S478 in mPER2) ^30–32^. We demonstrate that the period shortening induced by the tau-like CK1δ1-R178Q mutant^34^ can be partially replicated by forcing nuclear accumulation of wild-type CK1δ1 using an NLS tag (mNG-NLS-CK1δ1). Thus, the elevated nuclear accumulation of tau-like kinase may, in addition to its altered substrate specificity^32,61^, contribute to the period-shortening phenotype observed in tau mutant animals^62,63^.

This study examines the subcellular localization and interplay between PER:CRY:CK1δ complexes and CRY1. In the prevailing circadian model, transcriptional feedback is driven initially by the nuclear accumulation of PER:CRY:CK1δ repressor complexes. This is followed in the late phase by sustained repression mediated by CRY1 alone, which persists for several hours after CK1δ-dependent degradation of PER proteins. However, evidence from mouse liver^59^, SCN slices^18^, and U2OS cells^43^ and this study indicates that a substantial fraction of PER2 remains cytoplasmic, even when CRY1 is present, which normally supports nuclear import of the PER:CRY complex. While several mechanisms underlying cytoplasmic localization of PER2 have been proposed^38,64^, no circadian function has yet been attributed to the cytosolic PER2 pool.

We show by transient transfection and overexpression that the distribution of PER2 between cytoplasm and nucleus depends on the expression ratio of PER2 and CRY1. DOX-induced transient overexpression was necessary to analyze how PER2 and CRY1 influence each other’s subcellular localization and dynamics while excluding contributions from endogenous PER1 and CRY2. Our data suggest that PER2 dimers and dimers bound by a single CRY1 molecule are predominantly cytosolic, whereas dimers bound by two CRY1 molecules are enriched in the nucleus (see Fig 1E). In the physiological context of a functional clock, PER1, CRY2, PER2, and CRY1 are rhythmically expressed with consecutive peaks at different CT’s^12,65^. Since their expression ratios and levels vary over the course of a circadian cycle^65^, the composition and saturation of PER complexes by CRYs depend on both their specific expression levels, subcellular localization and binding affinities. Furthermore, our data show that CK1δ-mediated phosphorylation promotes dissociation of bound CRY1, thereby supporting the cytoplasmic accumulation of PER2 dimers containing one or no CRY1 molecule. In the context of a functional circadian clock in U2OS cells, we observe that the subcellular distribution of PER2 shifts from predominantly nuclear to increasingly cytoplasmic over the course of the circadian cycle, consistent with our overexpression data. Inhibition of CK1δ/ε reverses this process, leading to dephosphorylation of cytosolic PER2 and its accumulation in the nucleus. Consistent with these observations, hyperphosphorylated PER2 is predominantly cytoplasmic in mouse liver^59^ and U2OS cells^43^.

Incorporating these findings into the current circadian framework, we propose that the transition between the early and late repressive phases involves a decline in nuclear PER2:CRY1:CK1δ complex levels, driven by both PER2 degradation and cytoplasmic relocalization. PER2 degradation and reduction of nuclear PER2:CRY1:CK1δ complexes by export below a critical threshold could provide a time window for CLOCK dephosphorylation and facilitate subsequent rebinding of CLOCK:BMAL1 to DNA. In addition, cytoplasmic PER2:CRY1:CK1δ complexes could act as a reservoir, gradually releasing CRY1 in a CK1δ-dependent manner to support repression of DNA-bound CLOCK:BMAL1, complementing de novo CRY1 synthesis, which peaks as PER2 levels fall. The contribution to repression by newly synthesized CRY1 and by CRY1 released from cytosolic PER2 could vary by cell type and physiological conditions. Supporting this, our data show that nuclear CRY1 levels are markedly higher and more sustained in CRY1-expressing U2OStx cells that overexpress and accumulate PER2 and CK1δ in the cytosol.

The fact that both PER1/2:CRY1/2:CK1δ complexes and CRY1 alone repress CLOCK:BMAL1 raises the question of whether the mammalian circadian clock gains a functional advantage from maintaining both repressive modes. Published data indicate that CRY1 is present and functionally active throughout the entire circadian cycle^18,66^. Although both PER and CRY proteins are rhythmically expressed, quantitative analyses reveal that PER protein levels are very low when CLOCK:BMAL1 activity reaches its circadian maximum, whereas CRY1 remains at ∼50% of its peak level^18,65^. Moreover, ChIP-seq analyses in mouse liver show that CRY1 is bound to circadian promoters not only during the late repressive phase (CT0-4), but also at CT8-12, when CLOCK:BMAL1 activity is maximal^12^. Finally, studies of endogenously tagged PER2-LUC demonstrated that PER2 oscillates around an elevated baseline in CRY1 knock-out U2OS cells and around a reduced baseline in CRY2 knock-out cells^66^. These findings suggest that in U2OS cells, CRY1 partially represses CLOCK:BMAL1 even during the circadian peak of transcription. Together, these observations support the view that CRY1 attenuates CLOCK:BMAL1 activity across the entire circadian cycle. Consequently, whereas PER1/2:CRY1/2:CK1δ complexes primarily regulate the troughs of circadian gene expression, CRY1 may act to limit the peak amplitude of CLOCK:BMAL1-dependent transcription, thereby enhancing the dynamic range and responsiveness of the clock during this phase.

In summary, our data suggest that cytoplasmic CK1δ is readily available to assemble in a one-to-one complex with PER proteins. After nuclear import of the PER:CRY:CK1δ complex, the bound kinase will equilibrate with the nuclear CK1δ pool. Because the nuclear CK1δ concentration is low, rebinding of CK1δ may not allow saturation of PERs in steady state. This provides a mechanistic explanation for why overexpression of nuclear CK1δ shortens circadian period length while overexpression of cytosolic CK1δ has no effect. Moreover, our overexpression data suggest that CK1δ-mediated phosphorylation disrupts PER2-CRY1 complexes, promoting the buildup of cytosolic PER2 complexes with substoichiometric CRY1 bound, while unbound CRY1 may gradually accumulate in the nucleus, a process analogous to nuclear import of newly synthesized CRY1. In the physiological context of a functional clock, the redistribution of CRY1 could contribute to establishing the poised promoter state with CLOCK:BMAL1.

Many of our conclusions are derived from, synchronized DOX-induced overexpression of tagged clock proteins combined with time-lapse live-cell microscopy. While this approach was necessary to capture mechanistic insights that would be technically challenging or impossible to resolve under physiological protein levels, it will require further validation with other methods and in other systems. Nonetheless, our findings align well with and recontextualize independent observations of clock protein levels, phosphorylation states, and subcellular localization in mouse liver^11,59^ and the SCN^18,46^, thereby contributing to our understanding of the repressive phase of the core circadian clock.

## Methods

### Plasmids

For cloning using restriction enzymes. Enzymes used for cloning were obtained from New England Biolabs (NEB). Restriction digestion, dephosphorylation of vector backbone and ligation was performed as recommended by NEB. Linear DNA was purified using NucleoSpin Gel and PCR Clean-up kit from Macherey-Nagel. Plasmids were extracted with GeneJET Plasmid Miniprep kit from Thermo Fisher Scientific. For cloning with overlapping regions (restriction enzyme-free cloning), enzymes (Q5 DNA polymerase, restriction enzyme DpnI and T4 DNA polymerase) were obtained from NEB. DpnI digestion and exonuclease reaction with T4 DNA polymerase was performed in NEBuffer 2.1. *E. coli* were transformed with exonuclease treated PCR product(s) to facilitate *in vivo* plasmid assembly via overlapping DNA ends. pcDNA4/TO-derived vectors were constructed by cloning the coding regions of human casein kinase 1 isoform delta (CK1δ), murine Period circadian protein homolog 2 (mPer2), and murine cryptochrome 1 (mCry1) downstream of the CMV-TetO2 promoter which allows for DOX-inducible protein expression in cells harboring the tetracycline-repressor cassette (tx; i.e., U2OStx) and constitutively high expression in cells without the tetracycline-repressor cassette (i.e., HEK293T). Mutations, fluorescent tags, and epitopes were introduced by via ligation-free overlap cloning. All plasmid DNA sequences were sequence-verified prior to use.

### Cell lines

T-REx-U2OS (U2OStx; Life Technologies) and HEK293T (ATCC) cells were maintained in Dulbecco’s Modified Eagle Medium (DMEM) supplemented with 10% Fetal Bovine Serum (FBS) and 1x Penicillin-Streptomycin (Pen-strep). Cell culture reagents were obtained from Life Technologies. Cells were grown and maintained at 37°C in a humidified incubator containing 5% CO2.

For plasmid transfection, U2OStx cells were transfected with Xfect Transfection Reagent (Takaka) following the manufacturer’s protocol whereas HEK293T cells were transfected with polyethylenimine (PEI, made in-house) following the conventional protocol. To generate stable U2OStx cell lines, basal U2OStx cells were seeded in a 24-well plate and grown to confluence overnight. Cells were transfected with pcDNA4/TO vectors containing the desired constructs. Stable transfectants were selected by growing cells to sub-confluence in complete media supplemented with 50 µg/mL hygromycin and 100 µg/mL zeocin (Invitrogen) over the course of two to three weeks. Resulting single cell colonies after selection were isolated for further experimentation as previously described. To induce protein overexpression, cells were treated with 10 ng/mL DOX.

### Immunofluorescence

Cells were seeded on 12mm #1.0 glass coverslips (VWR, CAT# 631-1577P) in 500 µL media. Whenever applicable, transient transfection and/or DOX induction was performed. Immunofluorescence was done 24 h post-induction. Cells were washed with PBS and fixed with 4% paraformaldehyde for 10 min at room temperature. Cells were permeabilized with PBS + 0.1% Triton X-100 for 10 min, then washed three times and blocked with PBS + 5% heat-inactivated FBS for 1 h. Cells were then treated with the following primary antibodies diluted in PBS + 3% BSA + 0.25% Tween20 for 1 h at 37°C: PER2 (in-house, 1:50, rabbit polyclonal), PER1 (in-house, 1:100, rabbit polyclonal), BMAL1 (in-house, 1:200, rabbit polyclonal)^67,68^, CK1δ (Abcam ab85320, 1:1000), FLAG (Sigma-Aldrich F3165, 1:200). This was followed by washing with PBS, and treatment with corresponding secondary antibody: goat anti-rabbit conjugated to AlexaFluor488 (Thermo Fisher Scientific A-11008, 1:500) or goat anti-mouse conjugated to AlexaFluor488 (Thermo Fisher Scientific A-11001, 1:500) diluted in PBS + 3% BSA + 0.25% Tween20 for 1 h at 37°C. Cells were again washed three times, then mounted onto glass slides with ProLong™ Glass Antifade Mountant with NucBlue (Thermo Fisher Scientific P36983) and sealed with nail polish.

### Fluorescence, time-lapse, and live-cell confocal microscopy

For microscopy of fluorescently-tagged proteins and immunofluorescence, a Nikon Ni-E microscope was used (Nikon Imaging Center, Heidelberg) at a 60X oil immersion objective and the corresponding filter sets for DAPI, FITC, and Cy5. Exposure times were optimized within each experiment to obtain comparable fluorescence intensities. For image acquisition, the central focal plane was determined and images were taken as at 0.3 µm z-steps to a total 11 steps. For time-lapse microscopy, an Incucyte S3 system (Sartorius) was used at a 20X objective using the built-in filter sets set for the green and red channel. Exposure times were optimized within each experiment to obtain comparable fluorescence intensities. Images were taken at a regular schedule every 4 h until the end of the experiment typically 48 h later. For time-lapse, live-cell, confocal microscopy, a Nikon A1R confocal microscope was used (Nikon Imaging Center, Heidelberg) with a built-in TokaiHit on-stage environmental chamber. To prevent autofluorescence, colorless DMEM supplemented with normocin was used. Additionally, 60X oil immersion objective and the corresponding laser and filter sets for DAPI, FITC, TRITC, and TD (brightfield) were used. Exposure times were optimized within each experiment to obtain comparable fluorescence intensities. For image acquisition, the central focal plane was determined and images were taken as at 1.5 µm z-steps to a total 5 steps. Images were taken every 5 min for 17 h. All resulting images were exported and digital image analysis was performed in FIJI.

### Quantitative PCR

Cells were grown in p24 plates and induced with DOX for 24 h. RNA was extracted using peqGOLD TriFast Reagent following manufacturer’s protocols. cDNA was reverse-transcribed from 500 ng total RNA extract with Maxima Reverse Transcriptase (Thermo Fisher Scientific) following manufacturer’s protocols. Quantitative PCR (qPCR) was performed on a StepOne™ Real-Time PCR System (Applied Biosystems, Themo Fisher Scientific, CAT# 4376598) using Maxima SYBR Green qPCR Master Mix (Thermo Fisher Scientific) following manufacturer’s protocols. The following primer pairs below were used for their respective targets: CK1-F: AACCAAACACCCTCAGCTCCAC and CK1-R: GCCCCAGCAGCTCCATCACCAT; GAPDH-F: TGCACCACCAACTGCTTAGC and GAPDH-R: ACAGTCTTCTGGGTGGCAGTG. Relative gene expression was normalized to GAPDH and calculated using the ΔΔC_T_ method. Further normalization to the DOX-induced U2OStx sample was done to determine fold induction of gene expression.

### Immunoblotting

Protein extraction was performed using an in-house lysis buffer (25□mM Tris-HCl, pH 8.0, 150□mM NaCl, 0.5% Triton X100, 2□mM EDTA, 1□mM NaF, freshly added protease inhibitors: 1 mM phenylmethylsulfonyl fluoride (PMSF), leupeptin (5 μg/ml), and pepstatin A (5 μg/ml)) according to the conventional protocol. Briefly, cells were scraped and sample was collected via centrifugation. The sample was incubated in ice-cold lysis buffer on ice for 10 min and then incubated in a sonicated water bath for 10 min. Samples were centrifuged at 16,000 x g, 4 °C, 15 min taking the supernatant. Protein concentration was quantified via Nanodrop (NP80, Implen). For immunoblot analysis, protein samples were analzyed by SDS-polyacrylamide gel electrophoresis and transferred to a nitrocellulose membrane (cytiva, Amersham Protran, 0.45 µm pore size) via semi-dry blotting. To control for protein loading, membranes were stained with Ponceau S. Membranes were typically incubated in primary antibody at 4°C overnight and in secondary antibody conjugated to horseradish peroxidase at room temperature for 2 h. The following primary antibodies were used: anti-FLAG (Sigma-Aldrich F3165, 1:5000, mouse monoclonal), anti-CK1δ (abcam AF12G4, 1:5000, mouse monoclonal), anti-V5 (Invitrogen R960-25, 1:5000, mouse monoclonal); and the following secondary antibody was used for luminescent detection: goat anti-mouse IgG (H+L)-HRP conjugate (Bio-Rad 170-6516, 1:10000). For decoration, membranes were incubated in decoration buffer (100 mM TrisHCl pH 8.5, 2.2 x 10-2 % (w/v) luminol, 3.3 x 10-3 % (w/v) coumaric acid, 9.0 x 10-3 % (v/v) H2O2) and exposed for a time series to X-ray films (Fujifilm). Films were then developed (Medical film processor from Konica Minolta). Densitometric analysis of immunoblots was done using the built-in module in FIJI. The results were measured and plotted onto GraphPad Prism for downstream analysis (i.e., plotting over time post-CHX chase).

### LMB treatment; CHX chase; MG132, PF670, CalA treatments

Unless otherwise stated, induction of protein expression was typically performed by addition of 10 ng/mL doxycycline (DOX; Thermo Fisher Scientific CAT# 631311) 24 h prior to downstream experiments. To inhibit CRM1-depdendent nuclear export, LMB was added to cells to a final concentration of 20 nM; an equivalent volume of methanol was added to cells as a negative control. To arrest protein translation for a cycloheximide (CHX) chase, CHX was added to cells to a final concentration of 10 µg/ml. Untreated samples were taken as the 0 min CHX sample. Samples were then taken at 15, 30, 45, 60, and 90 min post-CHX for protein extraction and immunoblotting. To inhibit the proteosome, cells were treated with 20 µM MG132 (Abmole Bioscience CAT# AMO-M1902) for 4 h prior to immunoblotting; 4 µM MG132 for 4 h prior to immunofluorescence. To inhibit CK1δ; cells were treated with 1 µM PF670462 (Tocris Biosciences CAT# 3316) for 4 h prior to immunoblotting and immunofluorescence. To inhibit phosphatases, cells were treated with 80 nM CalA (LC Labs Cat# C-3987) for 1 h. All chemical treatments were performed with the corresponding solvent (vehicle) control: DMSO for MG132 and CalA, and H2O for PF670.

### *in vitro* phosphorylation assay and co-immunoprecipitation

HEK293T cells were transfected with V5-PER2 and CRY1-FLAG and the lysate was used as a starting material for an *in vitro* phosphorylation assay. CK1δΔtail was purified and used as the kinase for an *in vitro* phosphorylation assay as previously described^34^. As a negative control, a reaction mix containing all of the reaction buffer components except for the MgCl_2_ was incubated with the sample in the absence of additional kinase. The in vitro phosphorylation reaction was carried out at 4°C for 6 h. The reaction was stopped using PF670 prior to subsequent co-IP. For co-IP, Protein G Sepharose 4 Fast Flow (cytiva, 17061802) beads were used following the manufacturer’s protocol. Briefly, a 1:1 slurry was prepared by repetitive washing and buffer exchange with 67% TBS. The beads were then incubated with anti-V5 antibody (Thermo Fisher Scientific, R960) at 4°C while rotating for 4 h and used directly. Resulting samples following an *in vitro* phosphorylation assay were loaded onto the beads and incubated at 4°C overnight. Supernatant was removed, and the beads were then washed thrice with cold 67% TBS and gently centrifuged at 300 x g, 4°C, for 30 s. Bound protein was eluted using 2x Laemmli buffer without β-mercaptoethanol and boiling at 95°C for 5 min. Corresponding “Input” and “IP” samples were taken and loaded onto a PAGE gel. For the IP sample, an 8x equivalent was loaded compared to Input.

### Dexamethasone treatment and synchronization

As an additional synchronization cue aside from media change, U2OStx cells were synchronized using dexamethasone (DEX, dissolved in ethanol) at a working concentration of 1 µM. At the designated timepoints, cells were treated with DEX for 20 min, followed by washing with 1x PBS. The medium was then replaced with fresh D10 marking the start of the time post-DEX synchronization. Cells were then harvested at 4, 8, 12, and 16 h post-DEX synchronization.

#### Subcellular fractionation

To separate the cytosolic and nuclear cellular fractions, the NE-PER™ Nuclear and Cytoplasmic Extraction Kit (Thermo Fisher Scientific, Cat #78833) was used following the manufacturer’s instructions.

### Luciferase reporter assay

Bioluminescence of luciferase reporter genes was performed as previously described^34^. Briefly, samples were measured with a plate reader (EnSpire from PerkinElmer) placed in a temperature-controlled incubator (CLF Plant Climatics E41L1C8). Cells were seeded in a white 96-well plate and transfected using Xfect with *Bmal1p*-luc vector^69^. Cells were synchronized via medium change using media supplemented with either PBS or DOX. The plate was sealed and bioluminescence was recorded at 30 min intervals at 37°C. Three independent experiments with four technical replicates per cell line and treatment were carried out. Resulting bioluminescence traces were detrended using a sliding 24h-time window for baseline subtraction and a sliding 3h-time window for smoothening. Circadian period was determined using the curve-fitting function to a non-linear damped sine wave function in GraphPad Prism 10.

### Statistical Analysis

Data were plotted on GraphPad Prism 10. To test for differences between two samples, student’s t-test was performed. To test for difference between three or more samples, ANOVA was performed. Statistical significance was defined by a p value of < 0.05. All data are presented as means□±□standard deviations (SD).

## Acknowledgements

We thank Alessia Ruggieri und Michael Knop for providing mKATE-2 and mNeonGreen templates, respectively. This work was supported by the Deutsche Forschungsgemeinschaft, TRR186.

## Ethics declarations

### Data Availability

The data supporting the findings of this study are available within the article and its supplementary information files. This study includes no data deposited in external repositories.

### Material Availability

Plasmids and cell lines generated in this study are available from the lead contact, Michael Brunner with a completed materials transfer agreement.

---

**Fig EV1.**
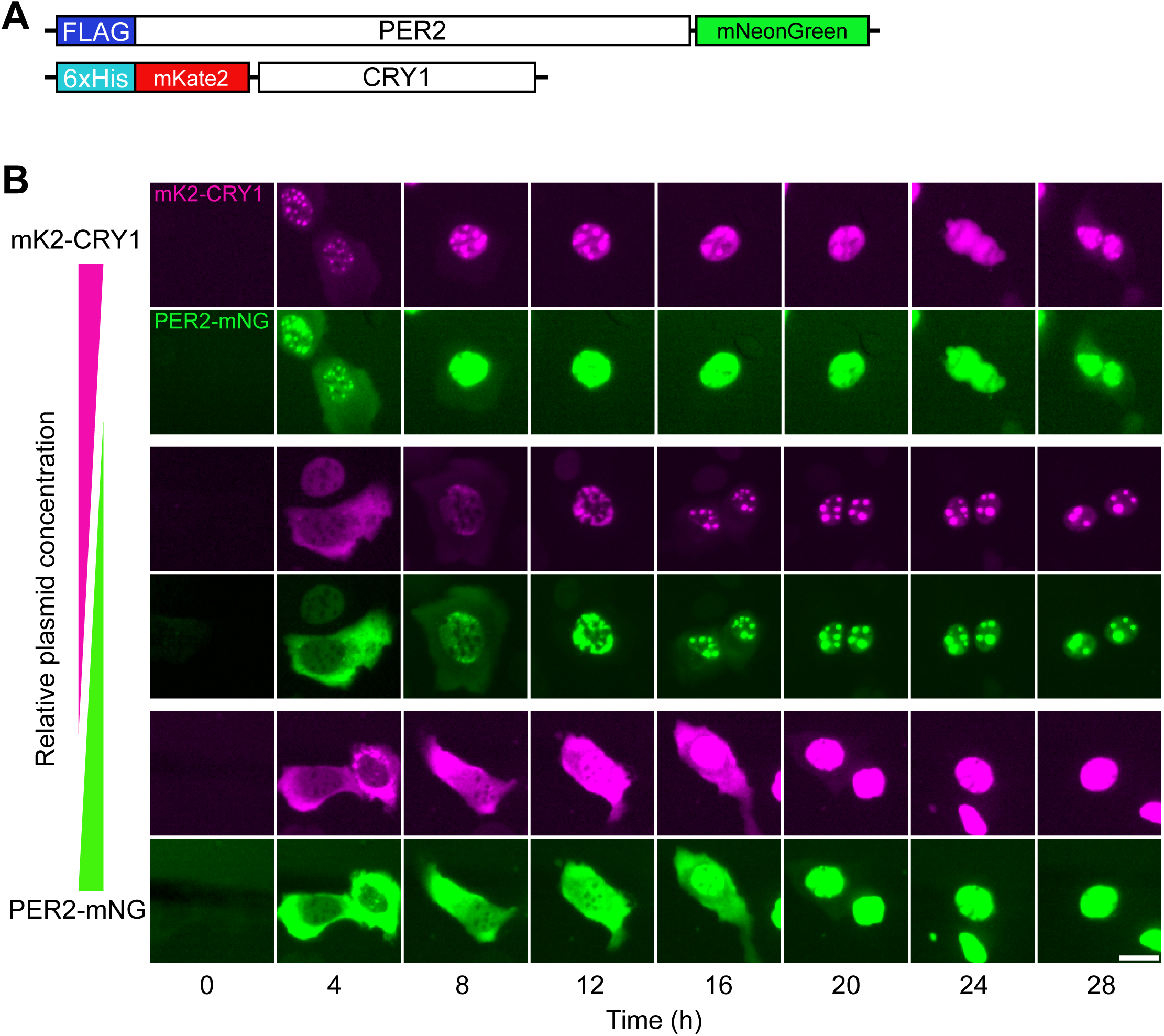
Subcellular dynamics of overexpressed PER2 and CRY1. (A) Constructs used in this study: FLAG–PER2–mNeonGreen (PER2–mNG) and 6×His–mKate2–CRY1 (mK2–CRY1). (B) Individual channels from the middle rows of Fig□1C, showing the localization of mK2–CRY1 (magenta) and PER2–mNG (green), respectively. Scale bar: 20□µm.

**Fig EV2.**
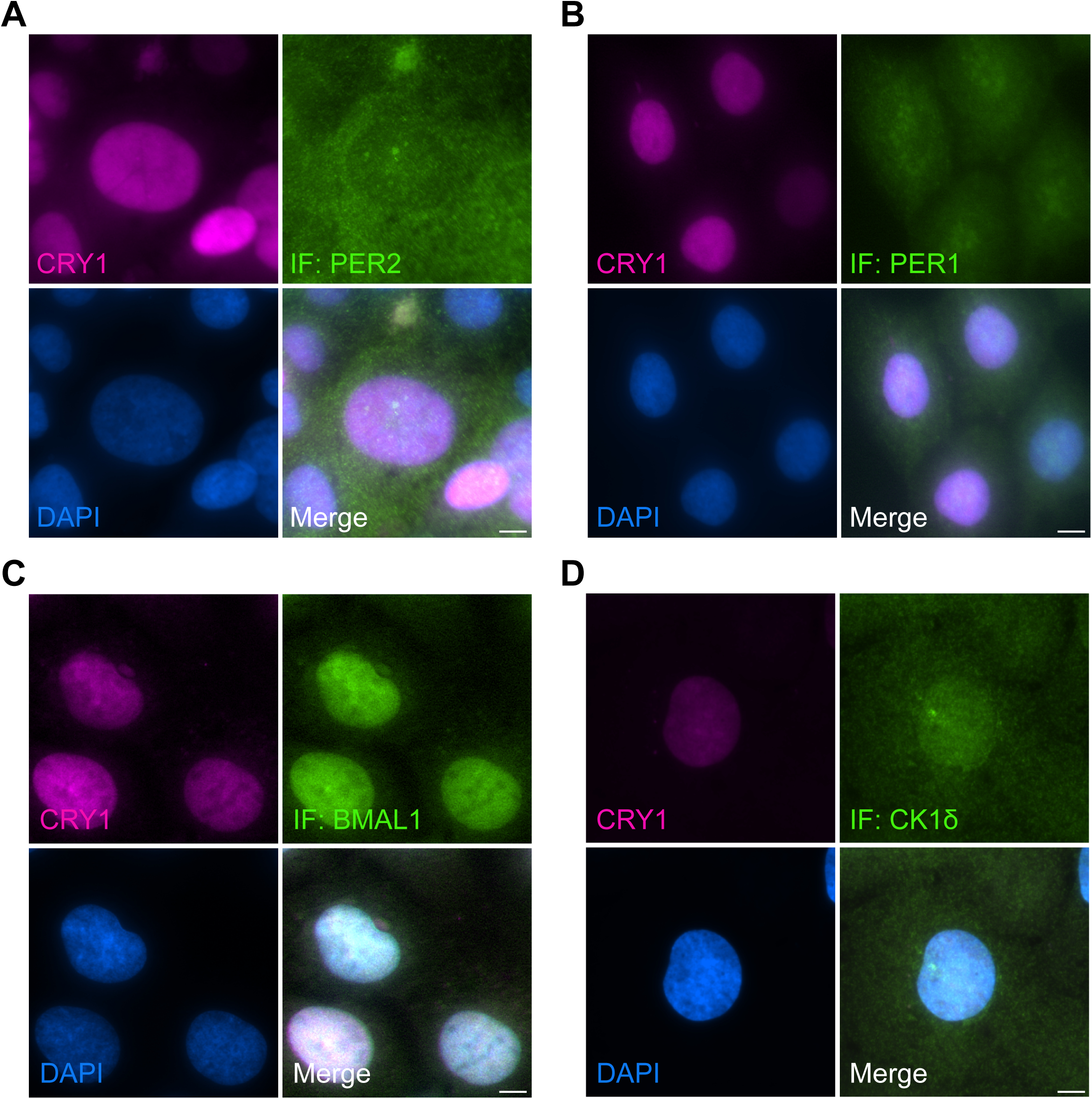
Analysis of untransfected U2OStx_mK2-CRY1 cells for control of PER2 transfected cells shown in Figure 2. Untransfected U2OStx_mK2-CRY1 cells displays predominantly homogeneous nuclear localization of mK2-CRY1 and (A) cytoplasmic and nuclear localization of endogenous PER2. (B) cytoplasmic and slightly enriched nuclear localization of endogenous PER1. (C) nuclear localization of endogenous BMAL1. (D) diffuse nuclear and cytoplasmic localization of endogenous CK1δ as well as localization at the centrosome. Scale bars 10 µm. n = 3.

**Fig EV3.**
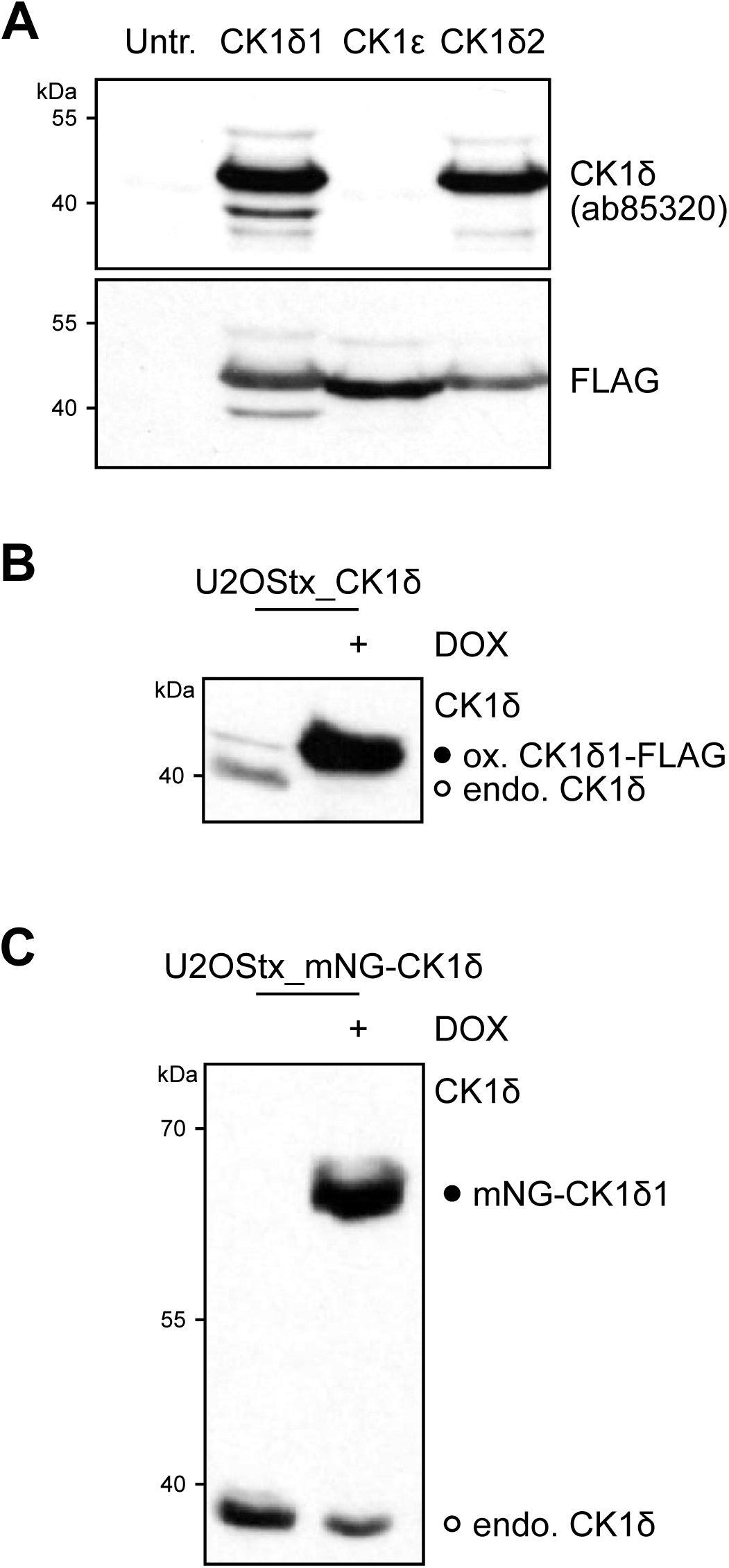
Endogenous CK1δ levels decrease upon overexpression of transgenic CK1δ. (A) CK1δ antibody (abcam, Cat# ab85320) recognizes CK1δ1 and CK1δ2 but not CK1ε. FLAG-tagged kinases were expressed in HEK293T cells. Untr.: untransfected control. Cell lysates were analyzed by Western-blot with antibody ab85320 (upper panel) and FLAG antibody (lower panel). n = 3. (B) Overexpression of CK1δ1-FLAG and (C) mNG-CK1δ1 reduce expression levels of endogenous CK1δ. Western blots with CK1δ antibody ab85320. n = 3.

**Fig EV4.**
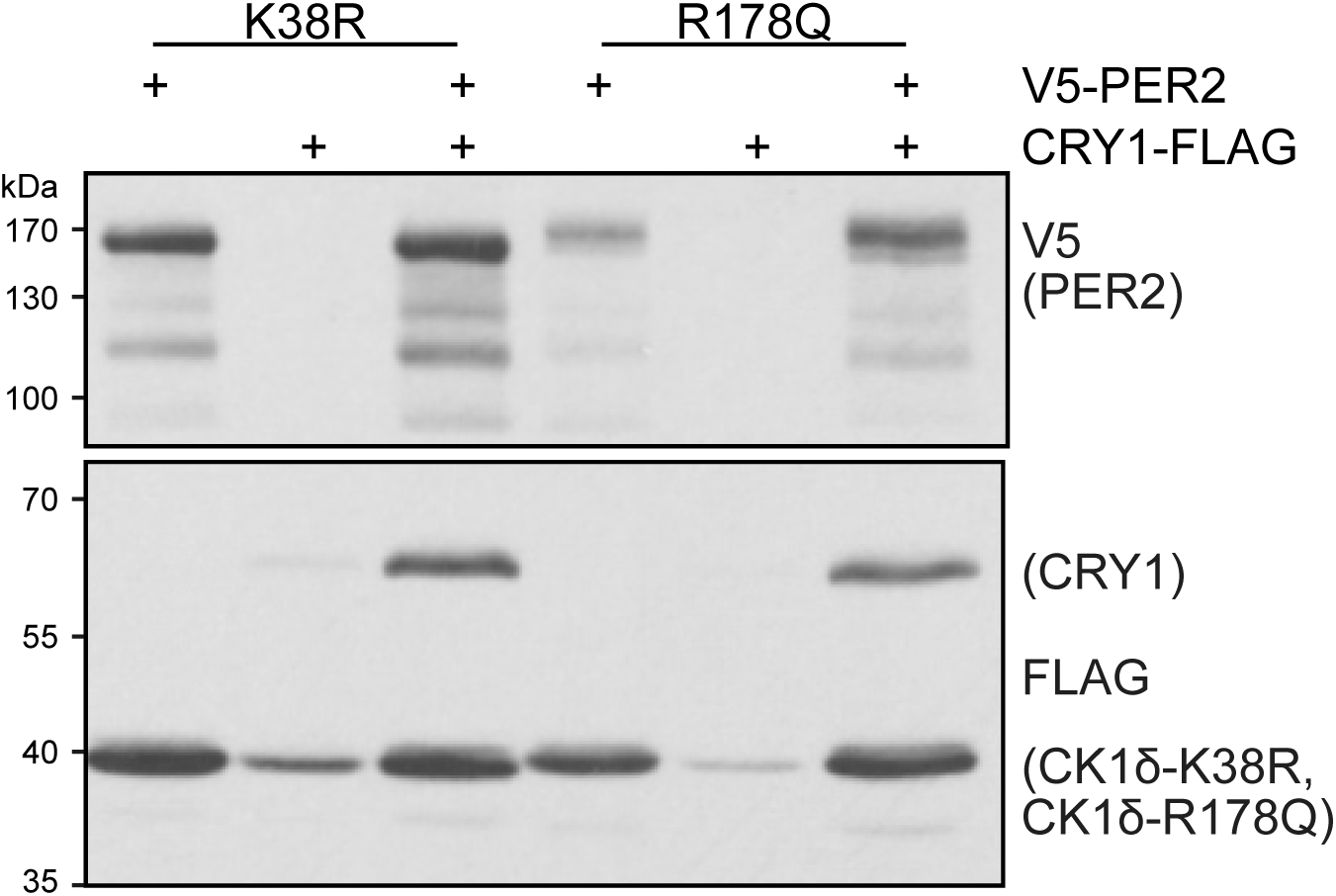
PER2 supports stabilization of mutant kinases. PER2 stabilizes CRY as well as CK1δ1-K38R and CK1δ1-R178Q, which are in absence of PER2 already more stable than CK1δ1 (see Fig 4c, from the same blot). HEK293T cell expression of indicated combinations of V5-PER2 and FLAG-tagged mK2CRY1, CK1δ1-K38R and CK1δ1-R178Q. n = 3.

**Fig EV5.**
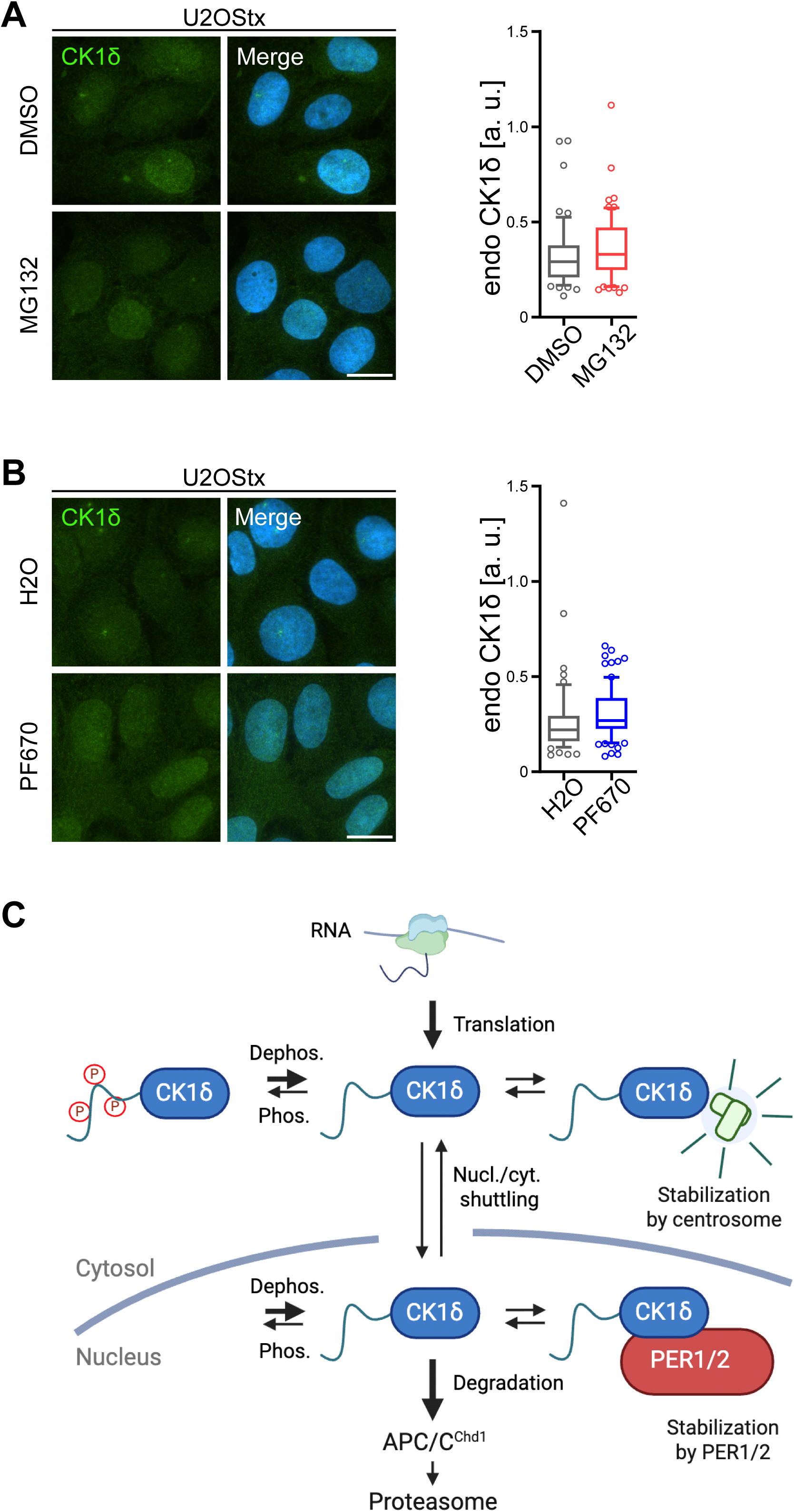
Impact of proteasome inhibition and kinase inhibition on endogenous CK1δ abundance and localization. (A) Proteasome inhibition by 5 µM MG132 for 4 h and (B) Kinase inhibition by 1 µM PF670 for 1 h does not result in accumulation of endogenous CK1δ. Scale bars 20 µm. Right figure parts: Quantifications of (A) and (B) show no significant change in abundance of endogenous CK1δ in inhibitor-treated cells compared to the vehicle control. n ≥ 50 cells, mean ± SD. (C) Schematic of CK1δ homeostasis. For details see main text.

**Fig EV6.**
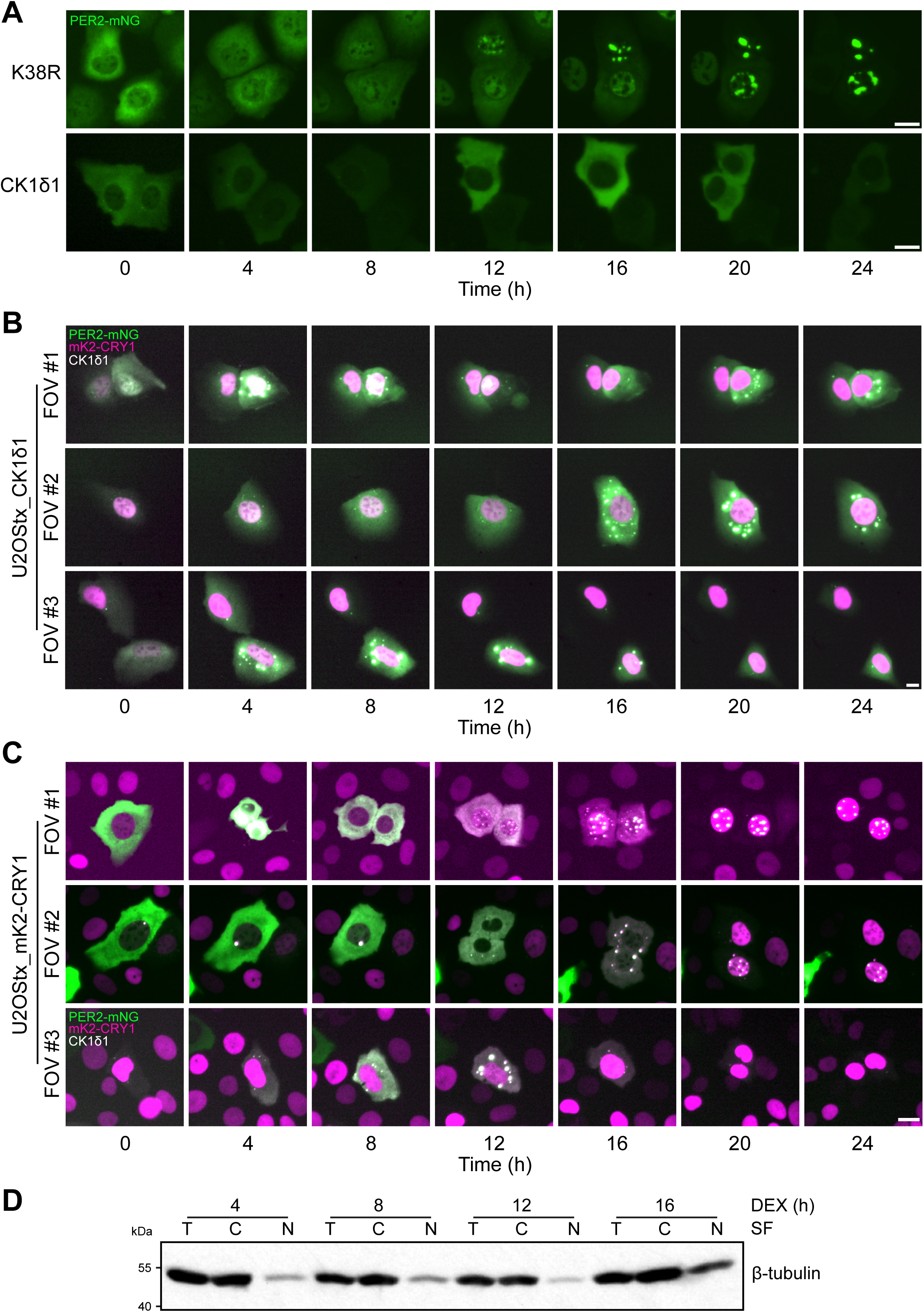
Active CK1δ promotes the formation of cytoplasmic PER2:CRY1 foci, serving as reservoirs for CRY1 release into the nucleus. (A) PER2-mNG alone accumulates slowly in the nucleus when expressed with inactive CK1δ-K38R and in the cytoplasm when expressed with active CK1δ. n = 3. (B) Relates to Fig 6A. Additional fields of view showing cytoplasmic PER:CRY foci in U2OStx_CK1δ1 cells transfected with PER2-mNG and mK2-CRY1. (C) Relates to Fig 6D. Additional fields of view showing that cytoplasmic PER2-CRY1 foci serve as a reservoir for the CK1δ1-dependent release of mK2-CRY1 into the nucleus after degradation of PER2-mNG. Scale bars 20 µm. (D) Relates to Fig 6F. β-tubulin was used as a loading and fractionation control.

**Fig EV7.**
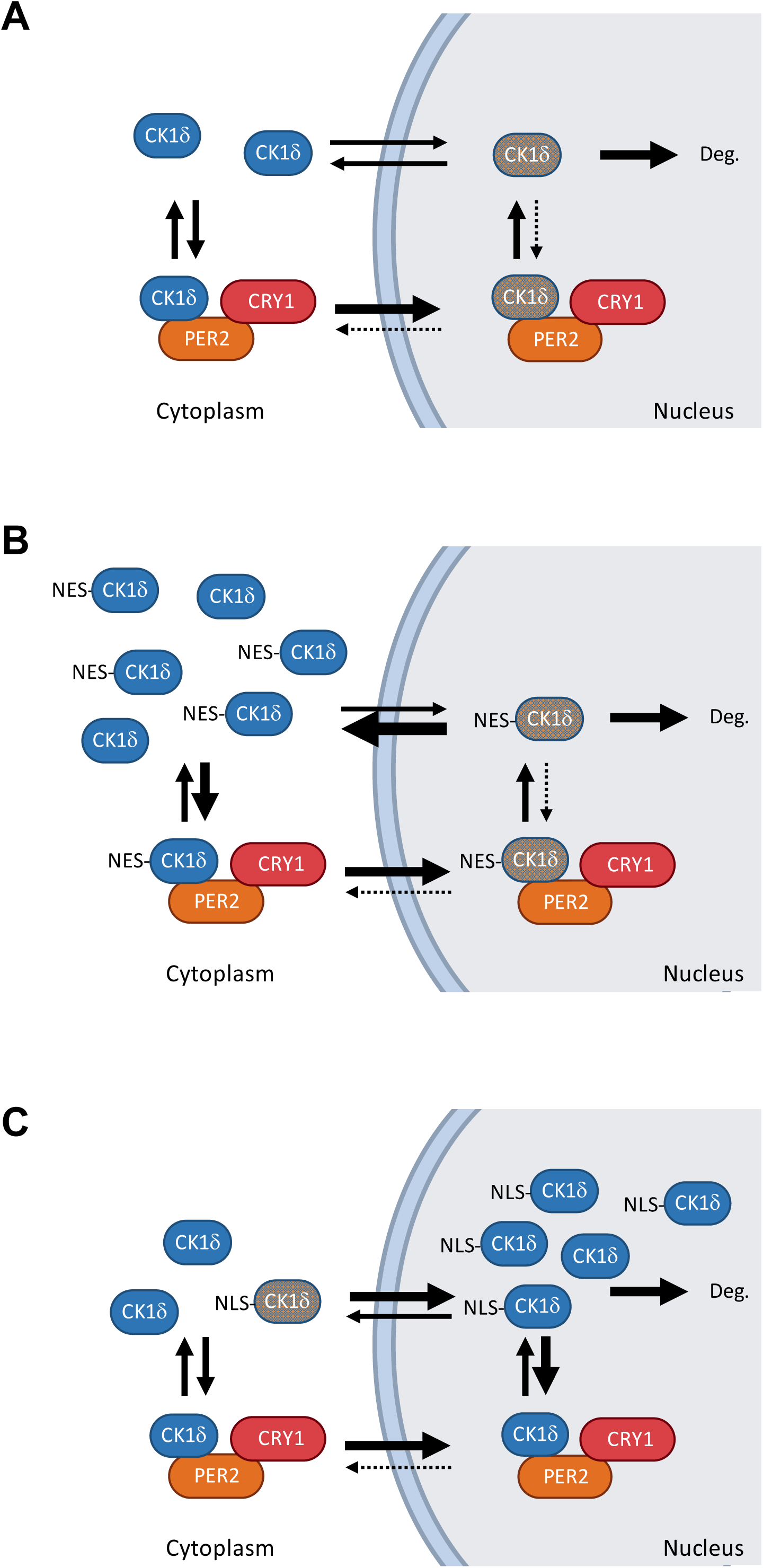
Model illustrating the impact of nuclear and cytoplasmic CK1δ on circadian period length. (A) Endogenous CK1δ continuously shuttles between the cytoplasm and the nucleus. Nuclear CK1δ levels remain low because the kinase is either rapidly degraded in the nucleus or efficiently exported back to the cytoplasm in a kinase activity-dependent manner. (B) Overexpression of NES-tagged CK1δ does not increase the nuclear kinase pool. The NES-tagged kinase enters the nucleus via PER:CRY complexes, with each PER molecule capable of importing only one NES-tagged kinase. Within the nucleus, PER-bound CK1δ equilibrates with the free nuclear pool. Once released from PER, rebinding of NES-tagged CK1δ is antagonized by either degradation or rapid export to the cytoplasm. Consequently, the nuclear kinase concentration remains low, and circadian period length is unaffected by NES-CK1δ overexpression. (C) Overexpression of NLS-tagged CK1δ increases the nuclear kinase pool by counteracting both nuclear export and degradation of CK1δ. This leads to a shortening of period length, as the elevated nuclear CK1δ facilitates PER2 binding and phosphorylation. Blue and gray schematic representations of CK1δ indicate high and low kinase concentrations, respectively.

